# Multimodal fusion of structural and functional brain imaging in depression using linked independent component analysis

**DOI:** 10.1101/676536

**Authors:** Luigi A. Maglanoc, Tobias Kaufmann, Rune Jonassen, Eva Hilland, Dani Beck, Nils Inge Landrø, Lars T. Westlye

**Author notes:** Corresponding authors: Luigi A. Maglanoc & Lars Westlye, Department of Psychology, University of Oslo, Pb. 1094, Blindern, 0317 OSLO, Norway.

## Abstract

**Background:** Previous structural and functional neuroimaging studies have implicated distributed brain regions and networks in depression. However, there are no robust imaging biomarkers that are specific to depression, which may be due to clinical heterogeneity and neurobiological complexity. A dimensional approach and fusion of imaging modalities may yield a more coherent view of the neuronal correlates of depression.

**Methods:** We used linked independent component analysis to fuse cortical macrostructure (thickness, area, gray matter density), white matter diffusion properties and resting-state fMRI default mode network amplitude in patients with a history of depression (n = 170) and controls (n = 71). We used univariate and machine learning approaches to assess the relationship between age, sex, case-control status, and symptom loads for depression and anxiety with the resulting brain components.

**Results:** Univariate analyses revealed strong associations between age and sex with mainly global but also regional specific brain components, with varying degrees of multimodal involvement. In contrast, there were no significant associations with case-control status, nor symptom loads for depression and anxiety with the brain components, nor any interaction effects with age and sex. Machine learning revealed low model performance for classifying patients from controls and predicting symptom loads for depression and anxiety, but high age prediction accuracy.

**Conclusion:** Multimodal fusion of brain imaging data alone may not be sufficient for dissecting the clinical and neurobiological heterogeneity of depression. Precise clinical stratification and methods for brain phenotyping at the individual level based on large training samples may be needed to parse the neuroanatomy of depression.

## Introduction

With an estimated prevalence of 4.4%, depression affects more than 300 million worldwide (World Health Organization, 2017) and is a substantial contributor to disability and health loss (Friedrich, 2017). Identifying useful imaging based and other biomarkers to aid detection of individuals at risk for depression and facilitating individualized treatment is a global aim (Cuthbert & Insel, 2012; Insel, 2014, 2015).

A host of studies across a range of neuroimaging modalities have implicated various brain regions and networks in depression. Meta-analyses of structural magnetic resonance imaging (MRI) studies have suggested thinner orbitofrontal (OFC) and anterior cingulate cortex (ACC) in patients with depression compared to healthy controls (Lai, 2013; Schmaal et al., 2017; Suh et al., 2019). A large-scale meta-analysis comprising 2148 patients and 7957 controls from 20 different cohorts reported slightly smaller hippocampal volumes in patients with depression compared to controls (Schmaal et al., 2016), but the overall pattern of results suggested substantial heterogeneity and otherwise striking similarity across groups for all other investigated subcortical structures (Fried & Kievit, 2016). A meta-analysis of diffusion tensor imaging (DTI) studies including 641 patients and 581 healthy controls reported fractional anisotropy (FA) reductions in the genu of the corpus callosum and the anterior limb of the internal capsule (Chen et al., 2016), implicating interhemispheric and frontal-striatal-thalamic connections among the neuronal correlates of depression. Supporting the relevance of brain connectivity in mood disorders, resting-state fMRI studies have reported aberrant connectivity within the default mode network (DMN) in patients with depression compared to healthy controls (Kaiser, Andrews-Hanna, Wager, & Pizzagalli, 2015; Mulders, van Eijndhoven, Schene, Beckmann, & Tendolkar, 2015; Yan et al., 2019).

However, despite meta-analytical evidence suggesting brain aberrations in large groups of patients with depression, the reported effect sizes are small and the direct clinical utility is unclear (Müller et al., 2017; Paulus & Thompson, 2019). One explanation for the lack of robust imaging-based markers in depression may be that previous studies have either focused on a single imaging modality or have analyzed different imaging modalities along separate pipelines and thus failed to model the common variance across features. In contrast, linked independent component analysis (LICA: Groves, Beckmann, Smith, & Woolrich, 2011; Groves et al., 2012) offers an integrated approach by fusing different structural and functional imaging modalities (Groves et al., 2012). LICA identifies modes of variation across modalities and disentangles independent sources of variation that may account both for large and small parts of the total variance, that may otherwise be overlooked by conventional approaches. By decomposing the imaging data into a set of independent components, LICA enables an integrated perspective that may improve clinical sensitivity compared to unimodal analyses (Alnæs et al., 2018; Doan, Engvig, Persson, et al., 2017; Francx et al., 2016; Wu et al., 2019).

Apart from the predominantly unimodal approaches in previous imaging studies, large individual differences and heterogeneity in the configuration and load of depressive symptoms represent other factors that could explain the lack of robust imaging markers. Symptom-based approaches have revealed more than 1000 unique symptom profiles among 3703 depressed outpatients based on only 12 questionnaire items (Fried & Nesse, 2015), suggesting large heterogeneity. Additionally, depression is highly comorbid with anxiety, with reported rates exceeding 50% (Johansson, Carlbring, Heedman, Paxling, & Andersson, 2013; Lamers et al., 2011). Furthermore, depression can be conceptualized along a continuum including individuals of the general, healthy population that may experience transient symptoms to varying degrees, and thus warrants a dimensional approach.

The main aim of the current study was to determine whether fusion of neuroimaging modalities would capture modes of brain variations which discriminate between patients with a history of depression (n = 170) and healthy controls with no history of depression (n = 71), and which are sensitive to current symptoms of depression and anxiety across groups. To this end, we used LICA to combine measures of cortical macrostructure (cortical surface area and thickness, and grey matter density), white matter diffusion properties (DTI-based FA, MD and RD), and resting-state fMRI DMN amplitude.

There is evidence of sex and age differences in the prevalence and clinical characteristics of depression, including lower age at onset of first major depressive episode in women compared to men (Marcus et al., 2005), and longer duration of illness and different symptoms in older compared to younger patients (Husain et al., 2005), which may reflect differential neuronal correlates. Therefore, we tested for main effects of age and sex and their interactions with the resulting brain components’ subject weights on group and symptoms.

In addition, we assessed the overall clinical sensitivity of all measures combined using machine learning to classify patients and controls and to predict symptom loads for depression and anxiety, which we compared with age prediction. Based on the above reviewed studies and current models we anticipated 1) that brain variance related to depression would be captured in components primarily reflecting the previously extended functional neuroanatomy of depression, including limbic and fronto-temporal networks and their connections. Irrespective of having a history of depression, we hypothesized 2) several strong age and sex differences, reflecting well documented age and sex-related variance in brain structure, including global thickness and volume reductions with increasing age, and larger brain volume and surface area in men compared to women. To the extent that having a history of depression interacts with sex and age-related processes in the brain, we hypothesized 3) interactions between the age-related trajectories and sex differences identified above with case-control status or symptoms of depression. To increase robustness and generalizability we corrected for multiple comparisons across all univariate analyses and performed cross-validation and robust model evaluation in the machine learning analyses.

## Materials and Methods

### Sample

Patients (n = 194) were primarily recruited from outpatient clinics, while healthy controls (n = 78) were recruited through posters, newspaper advertisements and social media. The patient group was drawn from two related clinical trials (ClinicalTrials.gov ID NCT0265862 and NCT02931487). All participants were evaluated with the Mini International Neuropsychiatric Interview (M.I.N.I 6.0: Sheehan et al., 1998). Exclusion criteria for all participants were MRI contraindications and a self-reported history of neurological disorders. The study was approved by the Regional Ethical Committee of South-Eastern Norway (REK Sør-Øst), and we obtained a signed informed consent from all the participants. Symptom loads for depression and anxiety were evaluated using the Becks Depression Inventory (BDI-II; Beck, 1996) and the Becks Anxiety Inventory (BAI; Beck & Steer, 1993) respectively. The demographics for the final sample (after exclusions, see below) are shown in table 1. The range of symptom load for depression and anxiety for the control group was from 0 to 20 and 0 to 19 respectively, while the range for the patient group was from 0 to 51 and 0 to 45 respectively (see Figure 1 for the distributions).

**Table 1.** Demographics of the final sample. 6 patients were missing information about ISCED level, 2 controls and 1 patient were missing Ham-D scores, 2 controls and 2 patients were missing AUDIT scores, and 2 controls and 4 patients were missing DUDIT scores. P denotes the p-value from group comparisons using Chi-Square test for sex, handedness, history of additional disorders, and current SSRI medication status while we used Mann-Whitney U tests for the rest.

|  | Controls (n = 70) | Patients (n = 171) | *p* |
| --- | --- | --- | --- |
| Sex (female, %) | 46 (66) | 120 (70) | 0.599 |
| Age (mean, SD) | 41.8 (13.1) | 38.7 (13.3) | 0.092 |
| Education level ISCED (mean) | 6.0 (1.0) | 5.9 (1.2) | 0.932 |
| Depression symptoms |  |  |  |
| BDI-II (mean, SD) | 1.6 (3.0) | 11.6 (10.4) | <0.001 |
| Anxiety symptoms |  |  |  |
| BAI (mean, SD) | 1.7 (2.8) | 8.1 (8.1) | <0.001 |
| Other |  |  |  |
| AUDIT (mean, SD) | 4.8 (3.3) | 6.3 (5.0) | 0.127 |
| DUDIT (mean, SD) | 0.5 (1.9) | 0.8 (2.5) | 0.123 |
| Left Handedness (N) | 7 | 6 | 0.087 |
| History of anxiety disorder (N) | 1 | 50 | <0.001 |
| History of (hypo)mania (N) | 0 | 23 | <0.001 |
| History of other Axis-I disorders (N) | 0 | 23 | <0.001 |
| No. of major depressive episodes (mean, SD) | 0 | 4.4 (5.9) | <0.001 |
| Currently medicated (SSRI, N) | 0 | 52 | <0.001 |

**Fig. 1.**
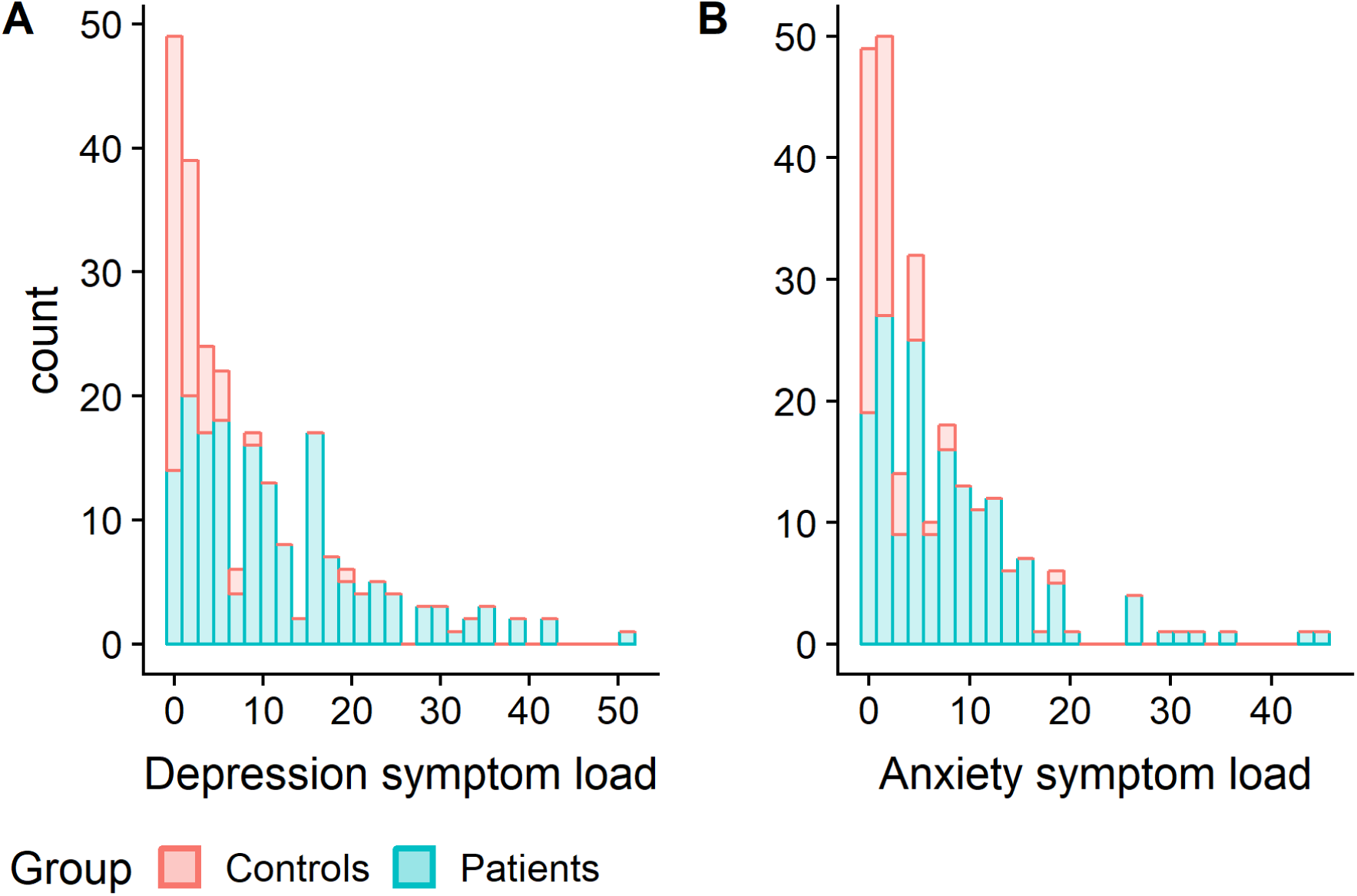
Histogram of symptom loads of (A) depression based on BDI-II and (B) anxiety based on BAI.

### Image Acquisition

MRI data was obtained on a 3T Philips Ingenia scanner (Phillips Healthcare) at the Oslo University Hospital using a 32-channel head coil. The same protocol was used for all participants, but there was a change in the phase-encoding direction during the course of study recruitment and data collection which affected the T1-weighted data for 4 controls and 95 patients, and resting-state fMRI data for 64 patients.

T1-weighted data was collected for 74 controls and 194 patients using a 3D turbo field echo (TFE) scan with SENSE using the following parameters: acceleration factor = 2; repetition time (TR)/echo time (TE)/ flip angle (FA): 3000 ms/3.61 ms/8°; scan duration: 3 min 16 s, 1 mm isotropic voxels.

Diffusion weighted data was collected for 72 controls and 184 patients using a dual spin echo, single-shot EPI sequence with the following parameters was used: TR/TE = 7200/86.5ms, FOV = 224 × 224 mm^2^, 112 × 112 matrix, 2.0mm isotropic voxels; 32 volumes with non-collinear directions (b = 1000s/mm^2^). Additionally, we acquired two b = 0 volumes with opposite phase polarity (blip up/down volumes).

Resting-state fMRI data was collected using a T2* weighted single-shot gradient echo EPI sequence was acquired for 72 controls and 178 patients with the following parameters: TR/TE/FA = 2500ms/30ms/80°; 3.00 mm isotropic voxels; 45 slices, 200 volumes; scan time ≈ 8.5 min. Participants were instructed to have their eyes open, and refrain from falling asleep.

### Structural MRI preprocessing

Vertex-wise cortical thickness and surface area measures (Dale, Fischl, & Sereno, 1999; Fischl, Sereno, & Dale, 1999) were estimated based on the T1-weighted scans using FreeSurfer (http://surfer.nmr.mgh.harvard.edu) (Fischl et al., 2002). Details are described elsewhere (Dale et al., 1999; Fischl et al., 1999) but in short, after gray/white boundary and pial reconstruction, cortical thickness was defined as the shortest distance between the surfaces vertex-wise (Dale et al., 1999), before resampling to the Freesurfer common template (fsaverage, 10,242 vertices; Fischl et al., 1999). The vertex-wise expansion or compression was used to calculate vertex-wise maps of arealization. None of the thickness nor surface area data for healthy control (n = 74) nor patients (n = 194) were excluded after visual QC.

### Voxel-based morphometry

Grey matter density maps (GMD) were created based on voxel-based morphometry (VBM) using the computational anatomy toolbox (CAT12: http://www.neuro.uni-jena.de/cat/) within SPM12 (http://www.fil.ion.ucl.ac.uk/spm/). This involved brain-extraction, gray matter-segmentation, and then registration to MNI152 standard space. The resulting images were averaged and flipped along the x-axis to create a left-right symmetric, study-specific grey matter template. The modulated gray matter maps were smoothed with a sigma of 4 mm (FWHM = 9.4 mm). None of the GMD data for healthy control (n = 74) nor patients (n = 194) were excluded after visual QC.

### DTI preprocessing

Processing steps included correction for motion and geometrical distortions based on the two b = 0 volumes and eddy currents by using FSL *topup* (http://fsl.fmrib.ox.ac.uk/fsl/fslwiki/TOPUP) and *eddy (*https://fsl.fmrib.ox.ac.uk/fsl/fslwiki/eddy). We also used *eddy* to automatically identify and replace slices with signal loss within an integrated framework using Gaussian process (Andersson & Sotiropoulos, 2016), which substantially improved the temporal signal-to-noise ratio (tSNR: Roalf et al., 2016) (*t* = 24.139, *p* < 0.001, Cohen’s d = 2.13). We fitted a diffusion tensor model using dtifit in FSL to generate maps of fractional anisotropy (FA), mean diffusivity (MD) and radial diffusivity (RD). Based on manual QC, we excluded 3 subjects due to insufficient brain coverage, and 3 subjects due to poor data quality. One additional subject was flagged with a tSNR of > 2 SD lower than the mean and discarded after additional manual QC. This yielded a total number of DTI scans of 71 healthy controls and 178 patients.

### Resting-state fMRI preprocessing

Resting-state fMRI data was processed using the FSL’s FMRI Expert Analysis Tool (FEAT). This included co-registration with T1 images, brain extraction, motion correction (MCFLIRT: Jenkinson, Bannister, Brady, & Smith, 2002), spatial smoothing (FWHM = 6 mm), high pass filtering (100s), standard space registration (MNI-152) with FLIRT, and single-session independent component analysis (ICA; MELODIC). Automatic classification and regression of noise components was done using ICA-based Xnoiseifier (FIX: Griffanti et al., 2014; Salimi-Khorshidi et al., 2014), with a threshold of 60. FIX substantially improved tSNR (*t* = 20.89, *p* < 0.001, Cohen’s d = 1.95), and no fMRI scans from healthy controls (n = 72) nor from patients (n = 178) were excluded. Group-level ICA with model order fixed at 40 was performed on a balanced subset of healthy controls and patients (N = 72 from each group), which has been used in a previous study (Maglanoc et al., 2019). Dual regression (Nickerson, Smith, Öngür, & Beckmann, 2017) was used to estimate spatial maps and corresponding time-series of all components. We then identified an IC representing the canonical DMN (Supplemental Figure 1) and used the individual DMN spatial maps from dual regression in multimodal decomposition using LICA.

### LICA

We used FMRIB’s LICA (http://fsl.fmrib.ox.ac.uk/fsl/fslwiki/FLICA) to perform data-driven multi-modal fusion, which evaluates shared inter-subject variations across the brain imaging measures (Groves et al., 2011, 2012). This produces spatial maps based on the commonalities across features (e.g. GMD, DTI measures, DMN maps) and subjects, and corresponding subject weights (i.e. the degree to which a subject contributes to a LICA component). We included complete data from 70 patients and 171 controls in the decomposition. We chose a relatively low model order of 40 based on previous recommendation of estimating robust components (Wolfers et al., 2017), and the biological meaningfulness of the spatial maps. For transparency and comparison, we also performed similar analysis using a higher dimensionality (80, more details in Supplemental). For both model orders, we discarded components which were highly driven by one subject (threshold: > 20%) yielding a total of 40 and 67 components respectively (Supplemental Figure 2). One component was strongly associated with phase encoding direction (IC4 in both decompositions, *t* = 33.07 and *t* = 32.47, *p* < 0.001 respectively, Supplemental Figures 3 and 4) but not removed from the analyses because of the biologically meaningful spatial patterns.

### Statistical analysis

Statistical analyses were performed in R version 3.5.1 (R Core Team, 2018) and Matlab 2014A (The MathWorks). We used linear models to test for main effects of clinical characteristics (case-control status, symptoms), age, and sex on each LICA subject weight with each IC as the dependent variable. In additional models we tested for interactions between age or sex and clinical characteristics (case-control status, symptoms) on each IC. For the analyses involving symptoms, one healthy control was removed due to missing data. We included phase encoding direction as an additional covariate in all the univariate analyses, and we controlled the false discovery rate (FDR) across tests using p.adjust in R.

### Machine learning approach

For group classification we submitted all LICA subject weights to shrinkage discriminant analysis (Ahdesmäki & Strimmer, 2010) in the R-package ‘sda’ (http://www.strimmerlab.org/software/sda/). For the main analyses we used the residuals of each component’s subject weight after regressing out age and sex, and additionally, phase encoding for IC4. As a supplemental analysis, we used the residuals of the subject weights after regressing out age, sex and phase encoding direction from all the ICs. For robustness and to reduce overfitting, we performed cross-validation with 10 folds across 100 iterations. We calculated area under the receiver operating curve (AUC) as our main measure of model performance using the R-package ‘pROC’ (Robin et al., 2011), but also accuracy, sensitivity and specificity. The relative feature importance was determined by calculating correlation-adjusted t-scores (CAT scores: Ahdesmäki & Strimmer, 2010). We determined statistical significance based on AUC using permutation-based testing across 10,000 iterations. We used the same framework to predict depression and anxiety symptoms, but by implementing shrinkage linear estimation (Schäfer & Strimmer, 2005) in the R-package ‘care’ (http://strimmerlab.org/software/care). Here, we computed root mean squared error (RMSE) between the raw and predicted scores as our main measure of model performance, but also mean absolute error (MAE), spearman’s rho, and R^2^. In this case, the relative feature importance was determined by computing the mean correlation-adjusted marginal correlation (CAR) scores (Zuber & Strimmer, 2011). Here, statistical significance was based on RMSE and permutation testing. As a comparison, we also predicted age using the same framework, using residuals of the subject weights after regressing out phase encoding direction in the methods shown above (using Pearson’s r instead of spearman’s rho).

## Results

### LICA

Figure 2A shows the degree of fusion across MRI measures for each component. Figure 2B shows the percentage of the total variance explained by each IC. Most of the components were characterized by region-specific features that were mainly bilateral, with the exception of 4 global components shown in Figure 3. Briefly, there was no substantial fusion between DMN maps and the other modalities, except for IC26 (Supplemental Figure 5).

**Fig 2.**
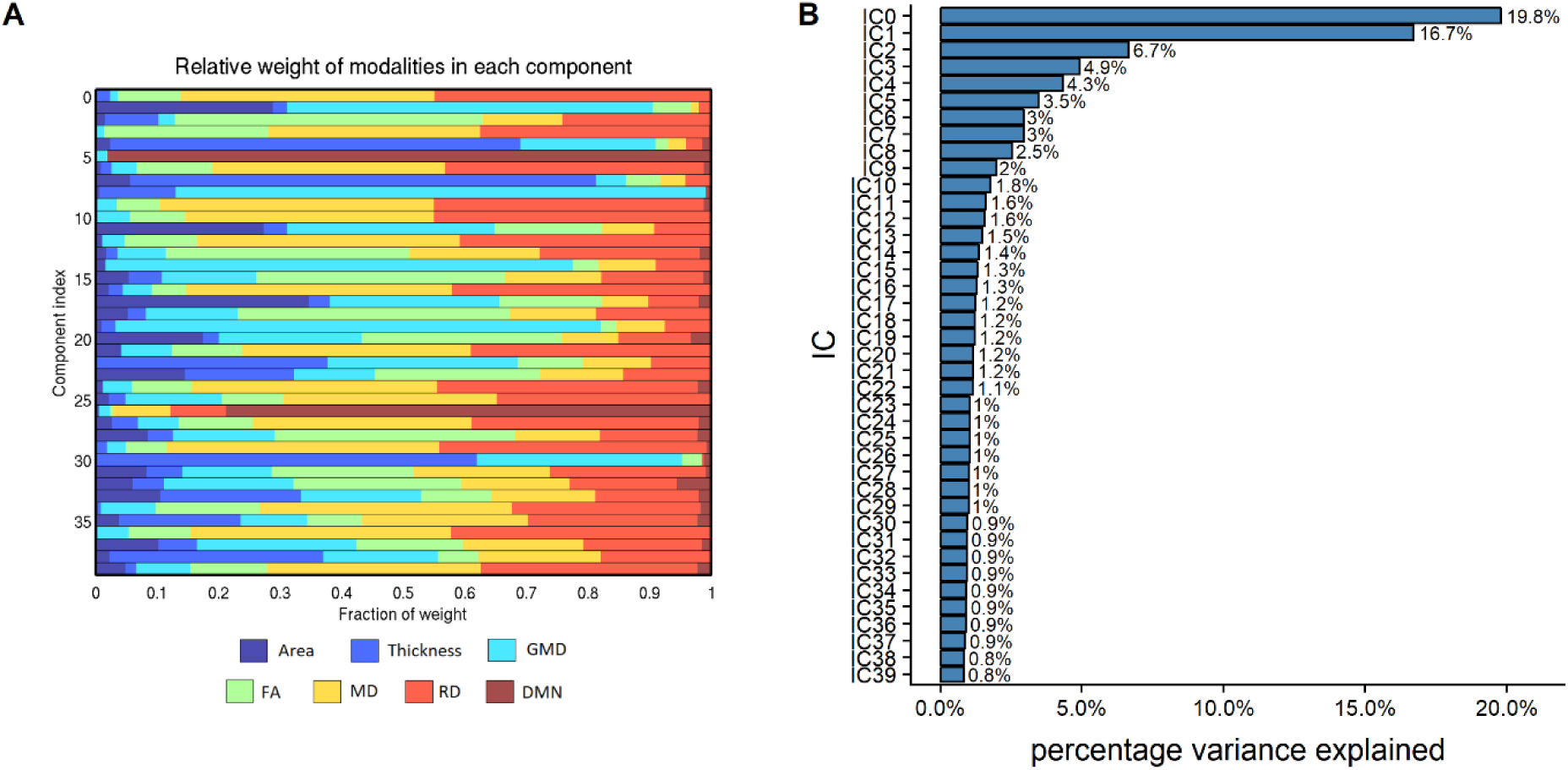
(A) The degree of fusing across MRI measures. (B) Explained percentage variance of each IC.

**Fig 3.**
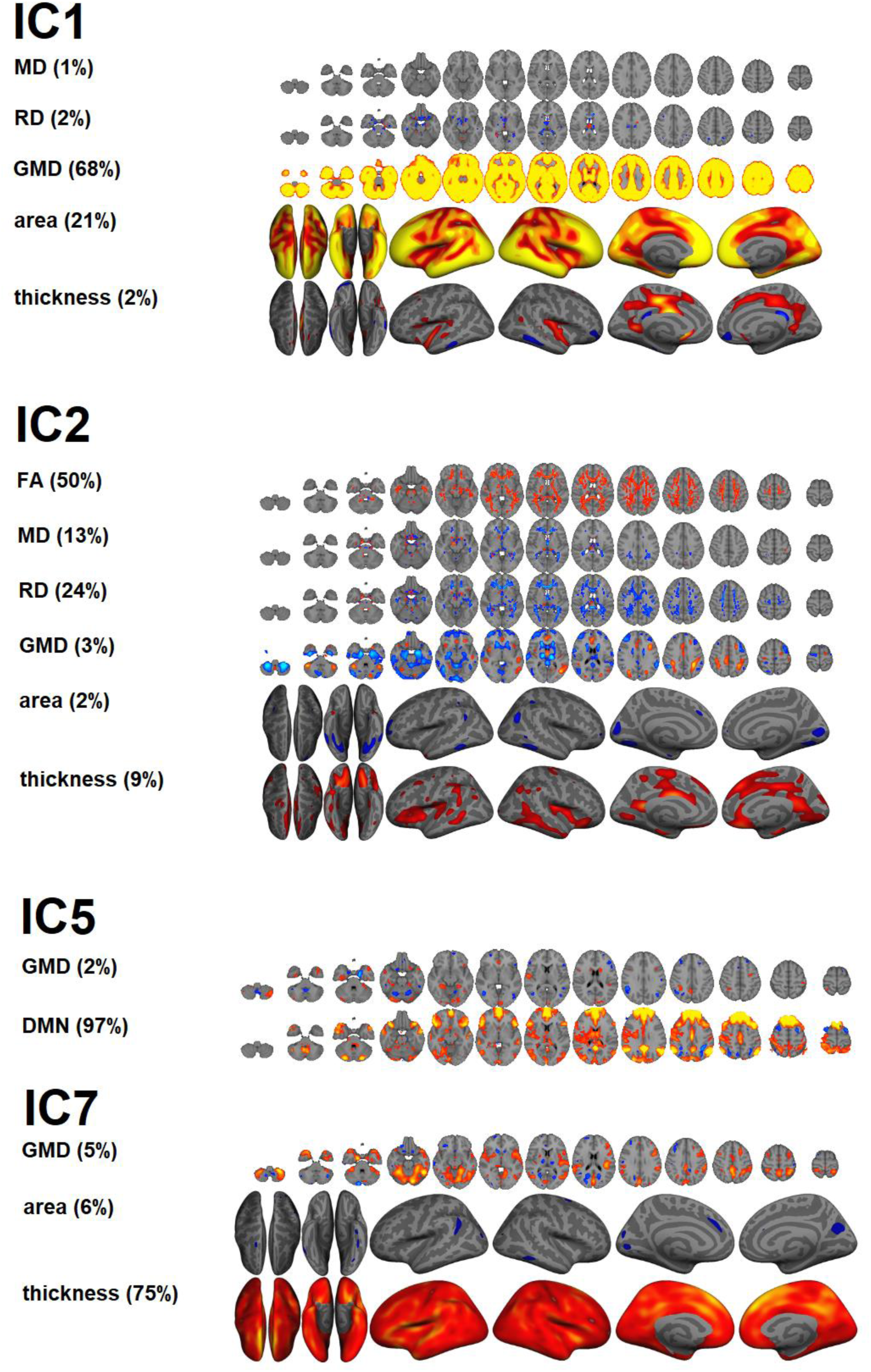
ICs that are mainly dominated by global features. For each IC, only measures that have an interpretable spatial pattern are presented. A z-score threshold of >= | 3 | was used for illustration. For visualization of the skeleton-based ICs, we used tbss_fill. IC1: global GMD and surface area. IC2: global white matter microstructure. IC5: DMN amplitude. IC7: global thickness.

### Univariate analyses

Table 2 shows results from linear models testing for main effects of group, age, sex, and symptom load for depression and anxiety on each IC. Supplemental Table 1 shows results from linear models testing for interactions between group or symptom loads for depression or anxiety with age or sex. Briefly, after corrections for multiple comparisons, the analysis revealed no significant associations between ICs and group, nor symptom load for depression and anxiety. There were significant main effects of age and sex on 10 and 5 LICA components (see Figure 4), respectively, but no significant main effects of group, with similar results for the decomposition with 80 components (see Supplemental Tables 2 and 3). Figure 3 shows the global LICA components associated with age and sex (see Supplemental Figure 6 for associations with additional LICA components). Age was negatively associated with IC1, indicating lower GMD and cortical surface area globally with increasing age, positively associated with IC2, indicating lower FA globally with increasing age, negatively associated with IC5, indicating lower DMN amplitude with increasing age, and negatively associated with IC7, indicating thinner cortex globally with increasing age. The analyses revealed main effects of sex on IC1, indicating larger global surface area and higher GMD in men compared to women, and IC5, indicating that men had higher DMN amplitude. The analyses revealed no significant interaction effects between group or symptom loads for depression or anxiety with age or sex with any of the ICs.

**Fig. 4.**
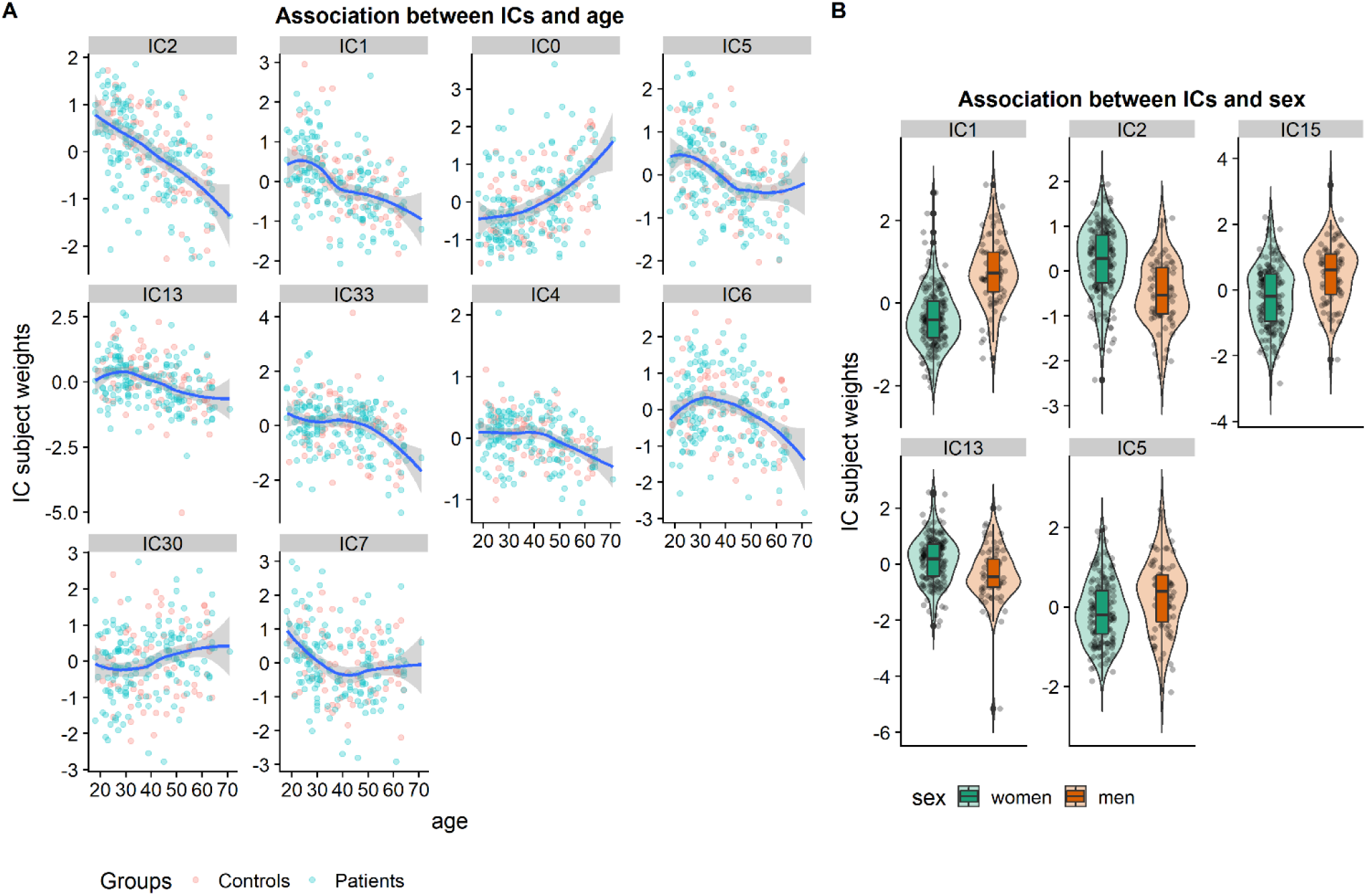
Significant effects of age and sex on ICs (*p* < 0.01). A: Scatter plots of the significant (linear) effects of age on ICs, sorted by the strength of the association. For visualization purposes we plotted LOESS and separated based on case-control status. The IC subject weights in the plot have been residualized for group, sex and phase encoding direction. B: Violin plots showing the distribution of the subject weights within men and women for each of the components showing a significant main effect of sex. The subject weights in the plot have been residualized for group, age and phase encoding direction

**Table 2.**
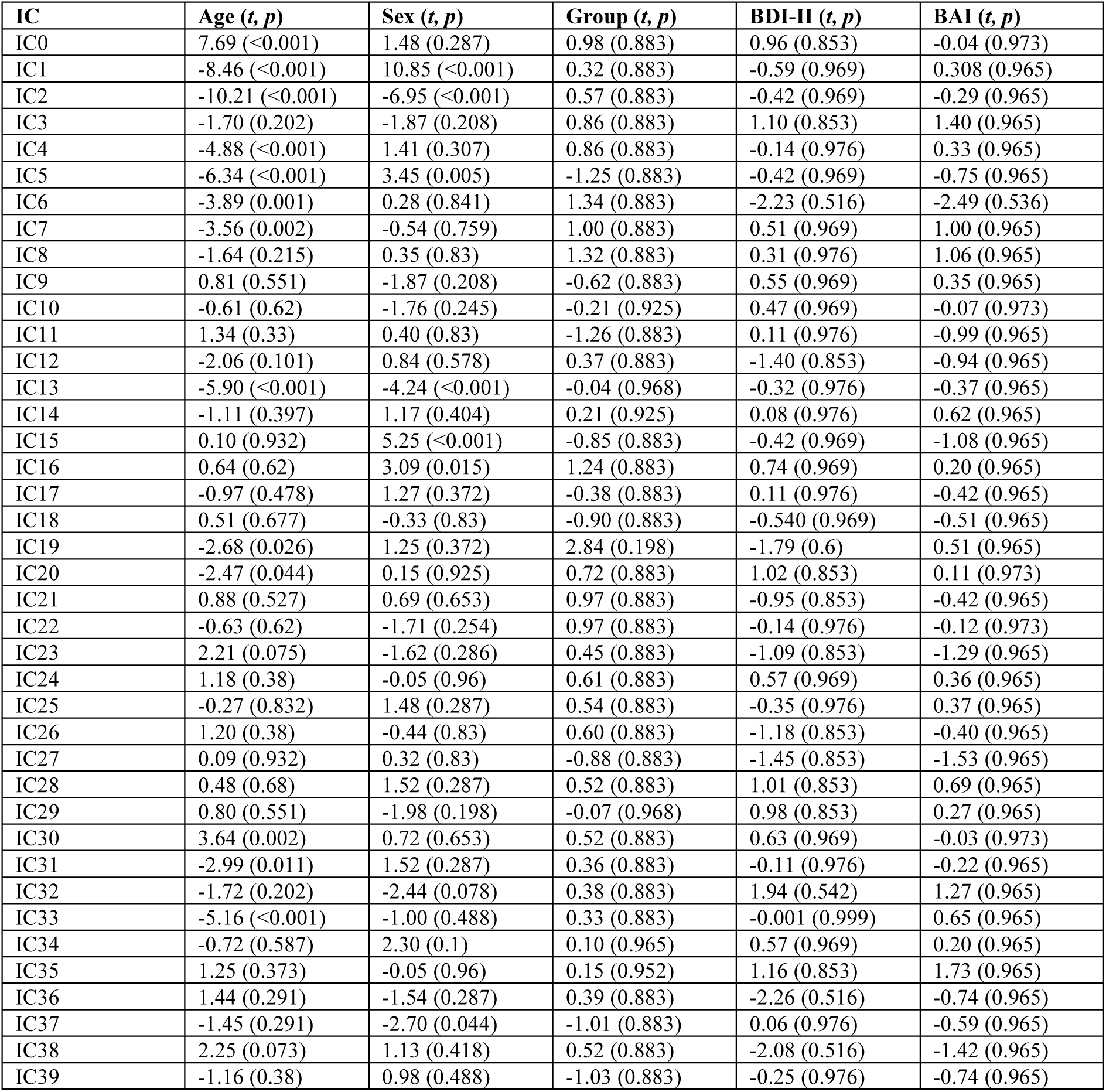
main effects of age, sex, group, symptom loads for depression and anxiety on ICs.

| IC | Age (*t, p*) | Sex (*t, p*) | Group (*t, p*) | BDI-II (*t, p*) | BAI (*t, p*) |
| --- | --- | --- | --- | --- | --- |
| IC0 | 7.69 (<0.001) | 1.48 (0.287) | 0.98 (0.883) | 0.96 (0.853) | -0.04 (0.973) |
| IC1 | -8.46 (<0.001) | 10.85 (<0.001) | 0.32 (0.883) | -0.59 (0.969) | 0.308 (0.965) |
| IC2 | -10.21 (<0.001) | -6.95 (<0.001) | 0.57 (0.883) | -0.42 (0.969) | -0.29 (0.965) |
| IC3 | -1.70 (0.202) | -1.87 (0.208) | 0.86 (0.883) | 1.10 (0.853) | 1.40 (0.965) |
| IC4 | -4.88 (<0.001) | 1.41 (0.307) | 0.86 (0.883) | -0.14 (0.976) | 0.33 (0.965) |
| IC5 | -6.34 (<0.001) | 3.45 (0.005) | -1.25 (0.883) | -0.42 (0.969) | -0.75 (0.965) |
| IC6 | -3.89 (0.001) | 0.28 (0.841) | 1.34 (0.883) | -2.23 (0.516) | -2.49 (0.536) |
| IC7 | -3.56 (0.002) | -0.54 (0.759) | 1.00 (0.883) | 0.51 (0.969) | 1.00 (0.965) |
| IC8 | -1.64 (0.215) | 0.35 (0.83) | 1.32 (0.883) | 0.31 (0.976) | 1.06 (0.965) |
| IC9 | 0.81 (0.551) | -1.87 (0.208) | -0.62 (0.883) | 0.55 (0.969) | 0.35 (0.965) |
| IC10 | -0.61 (0.62) | -1.76 (0.245) | -0.21 (0.925) | 0.47 (0.969) | -0.07 (0.973) |
| IC11 | 1.34 (0.33) | 0.40 (0.83) | -1.26 (0.883) | 0.11 (0.976) | -0.99 (0.965) |
| IC12 | -2.06 (0.101) | 0.84 (0.578) | 0.37 (0.883) | -1.40 (0.853) | -0.94 (0.965) |
| IC13 | -5.90 (<0.001) | -4.24 (<0.001) | -0.04 (0.968) | -0.32 (0.976) | -0.37 (0.965) |
| IC14 | -1.11 (0.397) | 1.17 (0.404) | 0.21 (0.925) | 0.08 (0.976) | 0.62 (0.965) |
| IC15 | 0.10 (0.932) | 5.25 (<0.001) | -0.85 (0.883) | -0.42 (0.969) | -1.08 (0.965) |
| IC16 | 0.64 (0.62) | 3.09 (0.015) | 1.24 (0.883) | 0.74 (0.969) | 0.20 (0.965) |
| IC17 | -0.97 (0.478) | 1.27 (0.372) | -0.38 (0.883) | 0.11 (0.976) | -0.42 (0.965) |
| IC18 | 0.51 (0.677) | -0.33 (0.83) | -0.90 (0.883) | -0.540 (0.969) | -0.51 (0.965) |
| IC19 | -2.68 (0.026) | 1.25 (0.372) | 2.84 (0.198) | -1.79 (0.6) | 0.51 (0.965) |
| IC20 | -2.47 (0.044) | 0.15 (0.925) | 0.72 (0.883) | 1.02 (0.853) | 0.11 (0.973) |
| IC21 | 0.88 (0.527) | 0.69 (0.653) | 0.97 (0.883) | -0.95 (0.853) | -0.42 (0.965) |
| IC22 | -0.63 (0.62) | -1.71 (0.254) | 0.97 (0.883) | -0.14 (0.976) | -0.12 (0.973) |
| IC23 | 2.21 (0.075) | -1.62 (0.286) | 0.45 (0.883) | -1.09 (0.853) | -1.29 (0.965) |
| IC24 | 1.18 (0.38) | -0.05 (0.96) | 0.61 (0.883) | 0.57 (0.969) | 0.36 (0.965) |
| IC25 | -0.27 (0.832) | 1.48 (0.287) | 0.54 (0.883) | -0.35 (0.976) | 0.37 (0.965) |
| IC26 | 1.20 (0.38) | -0.44 (0.83) | 0.60 (0.883) | -1.18 (0.853) | -0.40 (0.965) |
| IC27 | 0.09 (0.932) | 0.32 (0.83) | -0.88 (0.883) | -1.45 (0.853) | -1.53 (0.965) |
| IC28 | 0.48 (0.68) | 1.52 (0.287) | 0.52 (0.883) | 1.01 (0.853) | 0.69 (0.965) |
| IC29 | 0.80 (0.551) | -1.98 (0.198) | -0.07 (0.968) | 0.98 (0.853) | 0.27 (0.965) |
| IC30 | 3.64 (0.002) | 0.72 (0.653) | 0.52 (0.883) | 0.63 (0.969) | -0.03 (0.973) |
| IC31 | -2.99 (0.011) | 1.52 (0.287) | 0.36 (0.883) | -0.11 (0.976) | -0.22 (0.965) |
| IC32 | -1.72 (0.202) | -2.44 (0.078) | 0.38 (0.883) | 1.94 (0.542) | 1.27 (0.965) |
| IC33 | -5.16 (<0.001) | -1.00 (0.488) | 0.33 (0.883) | -0.001 (0.999) | 0.65 (0.965) |
| IC34 | -0.72 (0.587) | 2.30 (0.1) | 0.10 (0.965) | 0.57 (0.969) | 0.20 (0.965) |
| IC35 | 1.25 (0.373) | -0.05 (0.96) | 0.15 (0.952) | 1.16 (0.853) | 1.73 (0.965) |
| IC36 | 1.44 (0.291) | -1.54 (0.287) | 0.39 (0.883) | -2.26 (0.516) | -0.74 (0.965) |
| IC37 | -1.45 (0.291) | -2.70 (0.044) | -1.01 (0.883) | 0.06 (0.976) | -0.59 (0.965) |
| IC38 | 2.25 (0.073) | 1.13 (0.418) | 0.52 (0.883) | -2.08 (0.516) | -1.42 (0.965) |
| IC39 | -1.16 (0.38) | 0.98 (0.488) | -1.03 (0.883) | -0.25 (0.976) | -0.74 (0.965) |

### Machine learning analyses

Figure 5 shows the results of the machine learning analyses, with the spatial maps of a select few top features for each model shown in Supplemental Figure 7. Model performance was low for classifying patients and controls using residualized IC features (AUC = 0.5702, *p* = 0.06135, accuracy = 0.6169, sensitivity = 0.4292, specificity = 0.3009). The feature importance based on CAT-scores identified IC19 as the most important feature for classifying group. IC19 represents a covarying pattern of high GMD in most cerebellar regions, low GMD in cerebellar crus II, and high GMD in the angular gyri.

Model performance was low for predicting depression symptoms using residualized IC features (RMSE = 10.72, *p* = 0.9236, MAE = 8.498, R^2^ = -0.3302, spearman’s rho = 0.009). IC0 had the highest feature importance based on CAR-scores (positive association). IC0 is characterized by a complex covarying pattern including low GMD in temporal regions, the thalamus and cingulate, and low thickness in the cingulate and fronto-temporal regions. In terms of white matter diffusion properties, IC0 is characterized by high FA in several pathways including the posterior thalamic radiation and low FA in the anterior thalamic radiation and fornix, with mostly the reverse pattern for MD and RD.

**Fig. 5.**
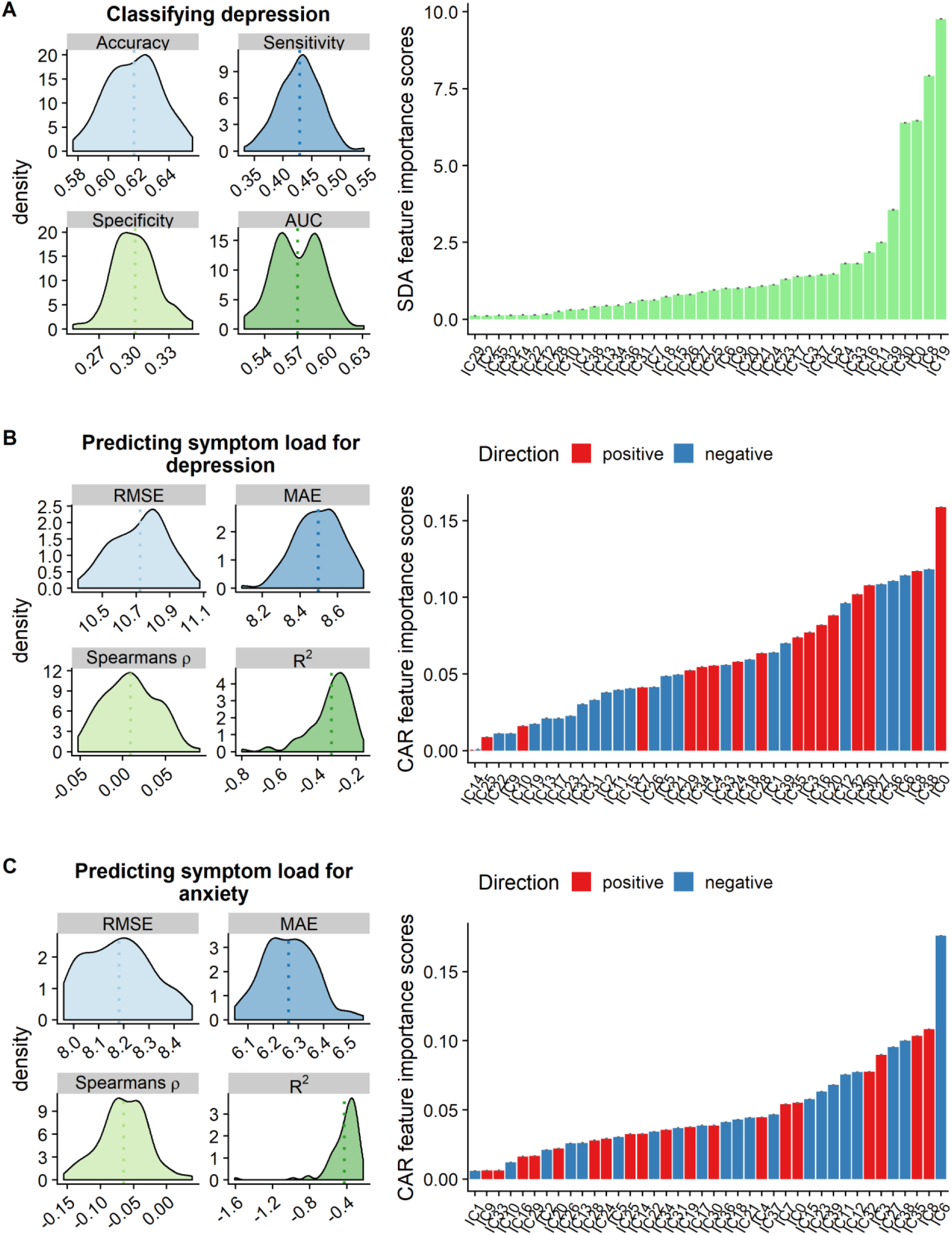
The results of the main analyses of the machine learning approach using 10-fold cross-validation across 100 repetitions for (A) classifying group (B) prediction symptom load for depression and (C) symptom load for anxiety. Here, phase encoding direction was only regressed out of the subject weights in IC4, while age and sex were regressed out from the subject weights of all the ICs. The figures on the left show prediction accuracy based on various model performance metrics. The barplots on the right show the most important features for each model based on CAT-scores (A) or CAR-scores (B and C).

Model performance was low when predicting anxiety symptoms using residualized IC features (RMSE = 8.181, *p* = 0.8946, MAE = 6.262, R^2^ = -0.424, Spearman’s rho = -0.064). IC6 had the highest feature importance based on CAR-scores (negative association). IC6 is mainly characterized by a complex covarying pattern of high FA in the splenium of the corpus callosum, high FA and low MD and RD in the fornix, high MD and RD in the thalamus, in addition to high GMD in the thalamus, and low GMD in hippocampal and amygdala regions. Model performance was slightly lower when regressing out phase encoding from all the IC features, and also suggested a different order of feature importance (see Supplemental Results and Supplemental Figure 8). Using the decomposition with higher model order revealed similar results in terms of feature importance, albeit slightly lower model performance for group classification, and symptom prediction (see Supplemental Results and Supplemental Figure 9). In contrast, model performance was high when predicting age (RMSE = 6.764, *p* < 0.0001, MAE = 5.530, R^2^ = 0.712, r = 0.861) using residualized features, with feature importance generally in line with the univariate results (see Figure 6). Model performance for predicting age was high but slightly lower when regressing out phase encoding from all the IC features (see Supplemental Results and Supplemental Figure 10), and when using the decomposition with higher model order (see Supplemental Results and Supplemental figure 11).

**Fig. 6.**
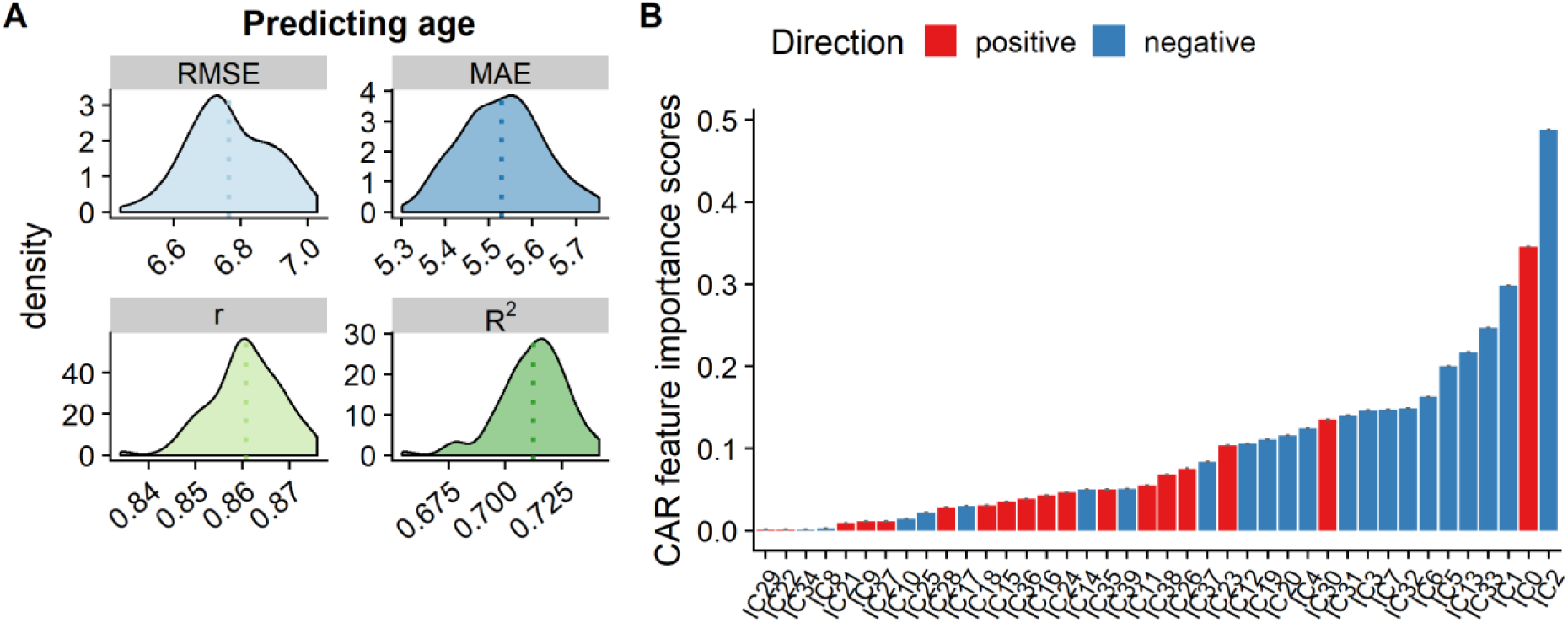
Age prediction regressing out phase encoding from the subject weights in IC4. (A) model performance results and (B) feature importance

## Discussion

The distributed functional and structural neuroanatomy of complex traits and disorders warrants integrated perspectives and analytical approaches. To this end, we probed the neuronal correlates of depression using multimodal fusion across cortical macrostructure, white matter diffusion properties and DMN amplitude based on resting-state fMRI. LICA yielded 40 components with various degree of multimodal involvement and different anatomical distributions, including both global and regionally specific patterns. Univariate analyses revealed strong associations with age and sex for several components’ subject weights after multiple comparison correction, but no robust group differences between patients with a history of depression and healthy controls, and no significant interactions between group and sex nor between group and age. Likewise, we observed no robust associations with symptom loads for depression or anxiety, nor interactions with age or sex. In line with the univariate analyses, the machine learning approach revealed overall low prediction accuracy for group classification and prediction of symptom loads for depression and anxiety, but high prediction accuracy for age.

Our univariate analyses revealed no main effects of history of depression or symptom load for depression on any of the LICA components. The he machine learning analyses here revealed overall low predictive value both for case-control status and symptoms of depression and anxiety, which is generally in line with the univariate analyses and an increasing body of literature suggesting small differences in brain structure between patients with MDD and healthy controls (Schmaal et al., 2017, 2016; Varoquaux, 2018; Wolfers, Buitelaar, Beckmann, Franke, & Marquand, 2015). While considering the overall low performance, the most important feature for classifying patients with a history of depression from healthy controls was a component encompassing covarying patterns of both high and low GMD in cerebellar regions (IC19). There is some evidence that depression is linked to cerebellar regions that communicate with networks related to cognitive and introspective processing (Depping, Schmitgen, Kubera, & Wolf, 2018). Cerebellar structural characteristics have recently been demonstrated to rank among the most sensitive brain features when comparing adult patients with schizophrenia and healthy controls (Moberget et al., 2018), and also for predicting psychiatric symptoms in youths (Moberget et al., 2019). The most important feature for predicting depression symptoms was IC0, which involves complex covarying patterns of low GMD and cortical thickness in mainly temporal but also frontal regions. This pattern is largely in line with previous research (Lai, 2013; Schmaal et al., 2017; Suh et al., 2019). IC0 also encompassed high FA and low MD and RD in interhemispheric connections and frontal-striatal thalamic pathways, in line with previous studies, albeit in the opposite direction (Chen et al., 2016). One study reported a positive association between symptom load for depression and FA in the thalamus (Osoba et al., 2013), and another study suggested this association (although in the opposite direction) is related to late onset MDD, especially in the corpus callosum (Cheng et al., 2014). However, while the implicated brain patterns are in line with previous reports, the overall poor model performance for predicting symptom load for depression warrants caution when interpreting this finding.

Higher age was related to lower global cortical thickness (IC7), in line with previous studies (e.g. Fjell et al., 2015). As hypothesized, higher age was also associated with lower global volume and smaller surface area (IC1), similar to previous studies using LICA (Doan, Engvig, Zaske, et al., 2017; Douaud et al., 2014), and a consistent finding in lifespan studies. Additionally, advancing age was negatively associated with IC2, indicating decreased FA globally, but also increased RD and to some extent MD, consistent with the aging literature (Davis et al., 2009; Sexton et al., 2014; Westlye et al., 2010). Higher age was associated with IC5, reflecting age-related decreases in DMN amplitude, in line with previous research (Damoiseaux et al., 2008; Mevel et al., 2013; Mowinckel, Espeseth, & Westlye, 2012; Razlighi et al., 2014; Vidal-Piñeiro et al., 2014). We observed high prediction accuracy for age in the machine learning approach, which shows the potential utility of LICA in estimating the gap between chronological and biological age (i.e. brain age gap).

Men had larger global brain volume and surface area than women (IC1), which is consistent with previous studies (e.g. Ritchie et al., 2018). Additionally, women had higher subject weights in IC13, reflecting lower FA in the corticospinal tract, portions of the superior longitudinal fasiculi and posterior thalamic radiation compared to men, generally in line with a large-scale UK Biobank study (Ritchie et al., 2018). We also found that men had greater DMN amplitude (IC5) than women, which adds to previous inconclusive findings (Mowinckel et al., 2012; Weissman-Fogel, Moayedi, Taylor, Pope, & Davis, 2010) and contrasts a previous report suggesting effects in the opposite direction (Jamadar et al., 2018).

In general, our analyses did not provide support for our hypothesis that a history of depression and symptoms of depression interact with age-related trajectories or sex differences of the LICA components. To the best of our knowledge, although studies have found specific cortical abnormalities related to adults with MDD, adolescents with MDD (Schmaal et al., 2017) and age at onset of depression (Ho et al., 2019), few or no studies have reported age-by-group interactions. Although there have been some early reports of a sex-by-group interaction in hippocampal volumes (e.g. Frodl et al., 2002), the recent large-scale ENIGMA MDD study reported no sex-by-group interactions in any subcortical volumes (Schmaal et al., 2016). Another recent study identified sex-by-group interactions using VBM, including higher GMD in the left cerebellum of male patients only, and lower GMD in the dorsal medial prefrontal gyrus in female patients only (Yang et al., 2017). However, the sample size was relatively small (less than 100 in the patient and control group each) which may affect the reproducibility of these findings. Despite separate reports of a link between resting-state DMN connectivity and rumination in depression (e.g. Hamilton, Farmer, Fogelman, & Gotlib, 2015) and evidence of sex-differences in rumination among adolescents (e.g. Jose & Brown, 2008), no studies have reported a sex-by-group interaction on DMN functional connectivity or amplitude.

The lack of positive findings in the univariate analyses and low predictive accuracy in the machine learning approach can be attributed to at least two factors. First, as illustrated by the large-scale ENIGMA studies (Ho et al., 2019; Schmaal et al., 2017, 2016), the effect sizes in neuroimaging studies of mental disorders and depression are overall small (Paulus & Thompson, 2019). Similarly, small sample sizes may contribute to over-inflated estimates of prediction accuracy in machine learning approaches (Wolfers et al., 2015). This is one possible explanation why other multimodal fusion studies of depression have reached higher prediction accuracies for classifying patients with depression from healthy controls (He et al., 2017; Ramezani et al., 2014; Yang et al., 2018), with patient groups consisting of no more than 60 individuals. Secondly, and also related to the small effect sizes, mental disorders including depression are clinically highly heterogenous. As an example, Müller and colleagues (2017) partially attribute the lack of convergence in their meta-analysis of activation-based fMRI experiments involving 1058 MDD patients to clinical heterogeneity. As a result, future research probing the neurobiology of depression should aim for large sample sizes (Rutledge, Chekroud, & Huys, 2019), and more importantly, stratifying patients (Feczko et al., 2019) with depression at the individual level. Pursuant to this, there has been considerable interest in identifying clinically relevant subgroups based on brain imaging, with initially encouraging results (Drysdale et al., 2017). However, the robustness and generalizability of such studies have been brought into question (Dinga et al., 2019), which may be partly due to substantial brain heterogeneity within groups, which has been illustrated in terms of morphometry in schizophrenia (Alnæs et al., 2019). Alternatively, dimensional measures such as brain age prediction (Kaufmann et al., 2018) and normative modelling (Marquand et al., 2019; Marquand, Rezek, Buitelaar, & Beckmann, 2016) have shown promising results in elucidating brain heterogeneity in mental disorders such as schizophrenia (Wolfers et al., 2018) and attention deficit/hyperactivity disorder (Wolfers et al., 2019).

The current findings should be considered in light of relevant limitations associated with statistical power and study design. The relatively low number of severely depressed patients may have influenced the sensitivity and specificity of the machine learning approach, in particular for predicting symptom loads of depression and anxiety. The varied current use of antidepressants in the patient group may have impacted the classification accuracy, although it is undoubtedly difficult to get large samples of non-medicated patients.

One weakness of this study is that the change in phase encoding direction may have introduced systematic differences in the MRI signal. However, only one component (IC4 in both decompositions) was strongly sensitive to phase encoding direction, suggesting that the remaining components were largely unaffected. Furthermore, we accounted for phase encoding direction in both the univariate and the machine learning analyses. This along with previous studies (Doan, Engvig, Persson, et al., 2017; Doan, Engvig, Zaske, et al., 2017; Groves et al., 2012) provides additional evidence that LICA is a promising tool to account for various scanner effects, particularly relevant for multi-site and longitudinal studies. We used a model order of 40 for LICA decomposition in the main analyses. Even though we did find similar feature importance ranking in the decomposition with higher model order, prediction accuracy was slightly lower across all models. Although there is no consensus on the optimal model order, this may imply that we were modelling more noise components in the higher model order decomposition, which was also supported by the number of discarded components due to dominance by a single subject.

In conclusion, based on fusion of structural, diffusion-weighted and resting-state fMRI data from 241 individuals with or without a history of depression, we identified multimodal and modality specific components that revealed strong associations with age and sex. None of the components showed significant association with categorical or dimensional measures of depression, nor any interaction effects with age and sex. Similarly, machine learning revealed low prediction accuracy for classifying patients from controls and predicting symptom loads. This study supports accumulating evidence of small effect sizes when comparing brain imaging features between patients with a history of depression and healthy controls, and indicates the need for more precise methods of stratifying individuals with depression, as well as large samples sizes.

## Acknowledgments

This work was supported by the South-Eastern Norway Regional Health Authority (2014097, 2015073, 2015052), the Research Council of Norway (249795, 229135), and the Department of Psychology, University of Oslo. The permutation testing was performed using resources provided by UNINETT Sigma2 - the National Infrastructure for High Performance Computing and Data Storage in Norway.

