## Supplementary material for "Multimodal fusion of structural and functional brain imaging in depression using linked independent component analysis": Supplmental Information

### *Supplemental Information*

#### Results

Machine learning results where age, sex and phase encoding direction have been regressed out from the subject weights of all the IC features.

*Case-control classification*

AUC = 0.5063, p = 0.848, accuracy = 0.5351, sensitivity = 0.3092, specificity = 0.3667.

*Predicting symptom loads of depression*

RMSE = 11.077, p = 0.9759, MAE = 8.801, $R^2$ = -0.4142, spearman's rho = -0.0994

*Predicting symptom loads of anxiety*

RMSE = 8.239, p = 0.9104, MAE = 6.334, $R^2$ = -0.445, Spearman's rho = -0.1022.

Machine learning results where phase age and sex have been regressed out from all the IC features, and phase encoding direction from IC4 at the higher decomposition (80, with a total of 67 components).

*Case-control classification*

AUC = 0.5385, Accuracy = 0.5813, Sensitivity = 0.37645, Specificity = 0.3317

*Predicting symptom loads of depression*

RMSE = 11.786, MAE = 9.180, $R^2$ = -0.6126, spearman's rho = 0.009

*Predicting symptom loads of anxiety*

RMSE = 9.235, MAE = 7.0402, $R^2$ = -0.8693, spearman's rho = -0.1019

Machine learning results for predicting age when regressing out phase encoding direction from the subject weights of all the ICs:

RMSE = 7.246, $p < 0.0001$, MAE = 5.988, $R^2 = 0.671$, $r = 0.838$

Machine learning results for predicting age when regressing out phase encoding direction from IC4 in the higher model order decomposition:

Age prediction: RMSE = 7.331, MAE = 5.942, $R^2 = 0.662$, $r = 0.838$

### Supplemental Tables

**Table S1**. Interaction effects of group, BDI and BAI with age and sex on ICs

| IC | Group x age (*t, p*) | Group x sex (*t, p*) | BDI x age (*t, p*) | BDI x sex (*t, p*) | BAI x age (*t, p*) | BAI x sex (*t, p*) |
|---|---|---|---|---|---|---|
| IC0 | -0.14 (0.947) | -0.22 (0.912) | -1.06 (0.899) | -1.10 (0.641) | 0.11 (0.976) | -0.82 (0.958) |
| IC1 | 0.31 (0.922) | 0.50 (0.823) | -0.34 (0.899) | 1.14 (0.641) | -0.55 (0.895) | 0.66 (0.958) |
| IC2 | 1.31 (0.813) | -0.17 (0.912) | 0.72 (0.899) | -1.11 (0.641) | 1.24 (0.895) | 0.30 (0.958) |
| IC3 | -0.40 (0.865) | 1.13 (0.823) | 0.36 (0.899) | 0.14 (0.982) | -1.24 (0.895) | -0.71 (0.958) |
| IC4 | 0.85 (0.813) | 1.05 (0.823) | 0.40 (0.899) | 0.61 (0.982) | 0.70 (0.895) | -0.15 (0.958) |
| IC5 | -0.65 (0.827) | -1.71 (0.587) | 0.44 (0.899) | -1.2 (0.641) | 1.08 (0.895) | 1.20 (0.788) |
| IC6 | 0.43 (0.865) | 0.56 (0.823) | 0.18 (0.956) | 0.58 (0.982) | -0.07 (0.976) | 0.27 (0.958) |
| IC7 | -0.79 (0.813) | -2.21 (0.379) | -0.47 (0.899) | -2.19 (0.479) | -0.90 (0.895) | -1.70 (0.609) |
| IC8 | 2.52 (0.167) | 2.20 (0.379) | 1.05 (0.899) | 0.24 (0.982) | 0.28 (0.976) | 0.20 (0.958) |
| IC9 | -0.79 (0.813) | 0.57 (0.823) | -0.49 (0.899) | 0.49 (0.982) | -0.76 (0.895) | -0.32 (0.958) |
| IC10 | 0.73 (0.816) | 0.67 (0.823) | 0.015 (0.988) | -0.16 (0.982) | 0.43 (0.895) | 0.34 (0.958) |
| IC11 | 0.82 (0.813) | 0.65 (0.823) | 0.29 (0.899) | 0.05 (0.982) | -0.03 (0.976) | -0.07 (0.958) |
| IC12 | -0.98 (0.813) | 1.42 (0.786) | -1.73 (0.703) | -1.12 (0.641) | -1.57 (0.895) | -1.74 (0.609) |
| IC13 | 0.76 (0.813) | 0.45 (0.823) | 0.84 (0.899) | 0.80 (0.942) | 1.48 (0.895) | 0.69 (0.958) |
| IC14 | 0.17 (0.947) | 0.46 (0.823) | 0.81 (0.899) | 0.26 (0.982) | 0.52 (0.895) | 0.24 (0.958) |
| IC15 | -1.88 (0.487) | 1.32 (0.823) | -0.57 (0.899) | 0.10 (0.982) | 0.06 (0.976) | -0.10 (0.958) |
| IC16 | 1.05 (0.813) | -0.17 (0.912) | 2.08 (0.703) | 0.10 (0.982) | 1.21 (0.895) | -0.42 (0.958) |
| IC17 | 2.08 (0.388) | 0.192 (0.912) | 0.06 (0.988) | -2.09 (0.479) | -0.64 (0.895) | -2.19 (0.609) |
| IC18 | 0.05 (0.962) | -0.08 (0.962) | 0.03 (0.988) | -1.39 (0.641) | -0.47 (0.895) | -1.13 (0.803) |
| IC19 | -2.53 (0.167) | 0.51 (0.823) | -0.39 (0.899) | -0.13 (0.982) | -0.71 (0.895) | -0.20 (0.958) |
| IC20 | 0.11 (0.947) | 0.17 (0.912) | 0.42 (0.899) | 0.24 (0.982) | 0.85 (0.895) | 0.48 (0.958) |
| IC21 | 2.53 (0.167) | 0.75 (0.823) | 1.63 (0.703) | -0.37 (0.982) | 0.45 (0.895) | -0.78 (0.958) |
| IC22 | 1.41 (0.813) | 1.72 (0.587) | 1.82 (0.703) | 1.58 (0.641) | 1.73 (0.895) | 0.60 (0.958) |
| IC23 | -1.32 (0.813) | -0.05 (0.963) | -1.63 (0.703) | -1.16 (0.641) | -2.20 (0.895) | -1.34 (0.729) |
| IC24 | 0.60 (0.827) | 2.59 (0.379) | 1.72 (0.703) | -0.20 (0.982) | 0.37 (0.921) | -0.21 (0.958) |
| IC25 | -1.20 (0.813) | 0.87 (0.823) | 0.33 (0.899) | 0.30 (0.982) | 0.65 (0.895) | 1.42 (0.729) |
| IC26 | -0.81 (0.813) | 0.76 (0.823) | 0.44 (0.899) | 1.99 (0.479) | -0.43 (0.895) | 0.56 (0.958) |
| IC27 | 1.06 (0.813) | -0.57 (0.823) | 0.47 (0.899) | -1.50 (0.641) | -0.64 (0.895) | -1.81 (0.609) |
| IC28 | -0.79 (0.813) | 0.90 (0.823) | 1.35 (0.871) | 1.68 (0.641) | 1.30 (0.895) | 1.95 (0.609) |
| IC29 | 0.27 (0.924) | 0.90 (0.823) | -0.40 (0.899) | -0.06 (0.982) | -0.81 (0.895) | 0.10 (0.958) |
| IC30 | 0.61 (0.827) | -0.80 (0.823) | -1.30 (0.871) | -1.39 (0.641) | -1.33 (0.895) | -2.17 (0.609) |
| IC31 | 1.15 (0.813) | 0.56 (0.823) | 0.60 (0.899) | 0.33 (0.982) | 0.07 (0.976) | -1.19 (0.788) |
| IC32 | -0.46 (0.865) | -0.49 (0.823) | 0.32 (0.899) | 0.13 (0.982) | -0.11 (0.976) | -1.44 (0.729) |
| IC33 | 0.10 (0.947) | 1.26 (0.823) | -0.27 (0.899) | 0.001 (0.999) | 0.03 (0.976) | -0.43 (0.958) |
| IC34 | 0.40 (0.865) | 0.41 (0.831) | 0.62 (0.899) | 1.14 (0.641) | 0.78 (0.895) | 0.48 (0.958) |
| IC35 | 0.59 (0.827) | 1.43 (0.786) | 0.49 (0.899) | 0.16 (0.982) | 0.16 (0.976) | -0.30 (0.958) |
| IC36 | 0.94 (0.813) | 1.80 (0.587) | 1.32 (0.871) | 2.14 (0.479) | 0.86 (0.895) | 0.89 (0.958) |
| IC37 | 1.14 (0.813) | -0.69 (0.823) | 0.10 (0.988) | -1.29 (0.641) | -0.51 (0.895) | -1.39 (0.729) |
| IC38 | -0.18 (0.947) | 0.44 (0.823) | -0.35 (0.899) | 0.15 (0.982) | -0.49 (0.895) | -0.83 (0.958) |
| IC39 | -0.46 (0.865) | -0.46 (0.823) | -1.06 (0.899) | 0.17 (0.982) | -0.47 (0.895) | 0.05 (0.958) |

**Table S2**. Main effects of age, sex, Group, BDI, and BAI on ICs from the higher model order decomposition

| IC | Age (*t, p*) | Sex (*t, p*) | Group (*t, p*) | BDI (*t, p*) | BAI (*t, p*) |
|---|---|---|---|---|---|
| IC0 | -7.54 (<0.001) | 8.92 (<0.001) | 0.47 (0.905) | -0.15 (0.962) | 0.56 (0.949) |
| IC1 | 7.29 (<0.001) | 0.78 (0.649) | 0.96 (0.905) | 1.24 (0.946) | -0.19 (0.949) |
| IC2 | -12.44 (<0.001) | -10.75 (<0.001) | 0.73 (0.905) | 0.09 (0.962) | -0.50 (0.949) |
| IC3 | 0.24 (0.889) | -4.09 (0.001) | 0.61 (0.905) | 0.65 (0.961) | 0.75 (0.949) |
| IC4 | -6.55 (<0.001) | 1.89 (0.252) | 1.30 (0.905) | 0.36 (0.962) | 0.40 (0.949) |
| IC8 | -4.81 (<0.001) | 2.23 (0.139) | 0.87 (0.905) | -1.26 (0.946) | -2.29 (0.767) |
| IC15 | -6.12 (<0.001) | 3.17 (0.018) | -1.19 (0.905) | -1.16 (0.946) | -0.89 (0.949) |
| IC16 | -1.12 (0.568) | 2.36 (0.108) | -0.63 (0.905) | 0.11 (0.962) | 0.19 (0.949) |
| IC17 | -1.03 (0.568) | -2.55 (0.069) | 0.63 (0.905) | 0.38 (0.962) | 1.04 (0.949) |
| IC20 | 1.57 (0.341) | -0.85 (0.649) | -0.15 (0.964) | 0.59 (0.961) | 0.39 (0.949) |

| | | | | | |
|---|---|---|---|---|---|
| IC22 | 1.19 (0.551) | -0.22 (0.906) | -1.13 (0.905) | -0.61 (0.961) | -0.55 (0.949) |
| IC23 | -0.98 (0.572) | -0.32 (0.865) | -0.29 (0.964) | 0.56 (0.962) | 0.17 (0.949) |
| IC25 | -3.16 (0.012) | 1.05 (0.55) | -0.81 (0.905) | -0.47 (0.962) | -0.36 (0.949) |
| IC26 | -2.25 (0.106) | 1.68 (0.289) | 2.35 (0.905) | -0.68 (0.961) | 0.55 (0.949) |
| IC27 | -1.05 (0.568) | 1.69 (0.289) | -0.18 (0.964) | 0.70 (0.961) | 0.56 (0.949) |
| IC28 | 6.01 (<0.001) | 0.52 (0.807) | 0.72 (0.905) | 1.31 (0.946) | 0.23 (0.949) |
| IC29 | 0.31 (0.86) | 0.16 (0.91) | -1.09 (0.905) | -0.85 (0.961) | -0.71 (0.949) |
| IC30 | -0.80 (0.607) | 3.14 (0.018) | -0.52 (0.905) | -1.49 (0.946) | -1.29 (0.949) |
| IC31 | 1.90 (0.195) | -3.93 (0.001) | 0.67 (0.905) | -0.08 (0.962) | -0.22 (0.949) |
| IC32 | -0.45 (0.781) | -0.42 (0.833) | 1.33 (0.905) | 0.18 (0.962) | -0.28 (0.949) |
| IC33 | 2.57 (0.052) | -1.54 (0.322) | 0.42 (0.905) | -1.08 (0.946) | -1.40 (0.949) |
| IC34 | -2.96 (0.02) | 5.29 (<0.001) | 1.44 (0.905) | -0.31 (0.962) | -0.10 (0.949) |
| IC35 | -0.91 (0.593) | 3.09 (0.019) | 1.50 (0.905) | 1.31 (0.946) | 0.26 (0.949) |
| IC36 | 0.83 (0.593) | -2.73 (0.051) | 1.65 (0.905) | 1.33 (0.946) | 1.05 (0.949) |
| IC37 | 0.15 (0.923) | 0.82 (0.649) | -0.72 (0.905) | -1.65 (0.946) | -1.52 (0.949) |
| IC38 | -0.50 (0.768) | 1.91 (0.252) | 0.75 (0.905) | 0.12 (0.962) | -0.24 (0.949) |
| IC39 | -1.09 (0.568) | -0.15 (0.91) | 0.23 (0.964) | 0.01 (0.994) | -0.10 (0.949) |
| IC40 | -4.80 (<0.001) | -0.38 (0.843) | -0.42 (0.905) | -0.27 (0.962) | 0.03 (0.988) |
| IC41 | 1.18 (0.551) | 1.84 (0.259) | 1.018 (0.905) | 0.71 (0.961) | 0.78 (0.949) |
| IC42 | 0.86 (0.593) | 1.09 (0.55) | -1.37 (0.905) | -1.05 (0.946) | -0.60 (0.949) |
| IC43 | 0.58 (0.727) | 1.55 (0.322) | 0.66 (0.905) | 2.18 (0.946) | 1.40 (0.949) |
| IC44 | -0.74 (0.632) | -1.26 (0.457) | 0.37 (0.92) | -0.36 (0.962) | -0.56 (0.949) |
| IC45 | -0.97 (0.572) | 0.71 (0.679) | -0.20 (0.964) | -1.06 (0.946) | -1.60 (0.949) |
| IC46 | -1.50 (0.362) | 0.22 (0.906) | -0.48 (0.905) | -0.54 (0.962) | -0.66 (0.949) |
| IC47 | 0.60 (0.718) | -1.77 (0.259) | 0.59 (0.905) | 0.21 (0.962) | 0.46 (0.949) |
| IC48 | -0.41 (0.787) | 1.46 (0.362) | -0.06 (0.987) | 0.23 (0.962) | 0.37 (0.949) |
| IC49 | 0.86 (0.593) | 0.59 (0.778) | -0.40 (0.905) | 0.71 (0.961) | -0.20 (0.949) |
| IC50 | -8.93 (<0.001) | -0.34 (0.864) | -0.02 (0.987) | 1.23 (0.946) | 2.13 (0.77) |
| IC51 | -2.78 (0.033) | 0.79 (0.649) | 0.52 (0.905) | 0.62 (0.961) | -0.15 (0.949) |
| IC52 | -1.79 (0.237) | 1.40 (0.391) | -0.26 (0.964) | -0.30 (0.962) | -0.39 (0.949) |
| IC53 | 1.25 (0.531) | -0.95 (0.597) | -0.04 (0.987) | 0.19 (0.962) | 0.71 (0.949) |
| IC54 | 1.95 (0.183) | -2.15 (0.155) | 0.44 (0.905) | -0.96 (0.961) | -0.43 (0.949) |
| IC55 | -0.42 (0.787) | 1.08 (0.55) | -0.02 (0.987) | 1.71 (0.946) | 0.94 (0.949) |
| IC56 | -2.10 (0.144) | 0.15 (0.91) | 1.82 (0.905) | 0.59 (0.961) | 0.57 (0.949) |
| IC57 | -0.04 (0.967) | 1.34 (0.418) | 1.49 (0.905) | 0.64 (0.961) | 0.33 (0.949) |
| IC58 | -0.76 (0.622) | 1.57 (0.322) | -0.42 (0.905) | -0.35 (0.962) | -0.74 (0.949) |
| IC59 | -1.64 (0.311) | 1.03 (0.55) | -0.89 (0.905) | -0.59 (0.961) | -0.83 (0.949) |
| IC60 | 0.84 (0.593) | -1.15 (0.525) | -0.13 (0.972) | 1.55 (0.946) | 0.92 (0.949) |
| IC61 | 0.52 (0.765) | -1.03 (0.55) | 1.10 (0.905) | 0.59 (0.961) | 0.24 (0.949) |
| IC62 | 0.47 (0.78) | -2.69 (0.051) | 1.45 (0.905) | -0.94 (0.961) | -0.89 (0.949) |
| IC63 | -0.89 (0.593) | -0.83 (0.649) | 0.59 (0.905) | 1.20 (0.946) | -0.11 (0.949) |
| IC64 | -1.45 (0.386) | 1.64 (0.299) | 0.25 (0.964) | 1.11 (0.946) | 0.92 (0.949) |
| IC65 | 0.05 (0.967) | 1.78 (0.259) | 0.23 (0.964) | -0.79 (0.961) | -1.72 (0.949) |
| IC66 | 1.06 (0.568) | -1.80 (0.259) | -0.17 (0.964) | -0.18 (0.962) | -0.50 (0.949) |
| IC67 | 2.03 (0.161) | -0.05 (0.971) | -1.11 (0.905) | -0.38 (0.962) | -0.41 (0.949) |
| IC68 | -0.14 (0.923) | 0.26 (0.904) | 0.53 (0.905) | 0.05 (0.977) | -0.02 (0.988) |
| IC69 | 0.23 (0.889) | 0.79 (0.649) | 1.13 (0.905) | 0.27 (0.962) | 0.14 (0.949) |
| IC70 | 0.99 (0.572) | 0.45 (0.825) | -1.13 (0.905) | -0.40 (0.962) | -0.44 (0.949) |
| IC71 | 1.03 (0.568) | -0.16 (0.91) | -0.52 (0.905) | -0.77 (0.961) | 0.23 (0.949) |
| IC72 | -1.04 (0.568) | 0.71 (0.679) | 0.05 (0.987) | 0.66 (0.961) | 0.79 (0.949) |
| IC73 | -2.34 (0.089) | -1.25 (0.457) | 0.81 (0.905) | 0.12 (0.962) | 0.88 (0.949) |
| IC74 | 1.52 (0.36) | -0.55 (0.796) | -0.75 (0.905) | -0.26 (0.962) | 0.36 (0.949) |
| IC75 | -2.57 (0.052) | 0.50 (0.808) | -0.98 (0.905) | 0.12 (0.962) | -0.31 (0.949) |
| IC76 | -0.22 (0.889) | 0.94 (0.597) | -0.70 (0.905) | -1.49 (0.946) | -0.95 (0.949) |
| IC77 | 0.90 (0.593) | -0.40 (0.843) | 0.82 (0.905) | -1.43 (0.946) | -0.34 (0.949) |
| IC78 | -0.65 (0.694) | -0.49 (0.808) | 0.78 (0.905) | 1.13 (0.946) | 2.35 (0.767) |
| IC79 | -0.13 (0.923) | -0.02 (0.985) | 0.82 (0.905) | 1.83 (0.946) | 0.56 (0.949) |

**Table S3**. Interaction effects of group, BDI and BAI with age and sex on ICs from the higher model order decomposition

| IC | Group x age (*t, p*) | Group x sex (*t, p*) | BDI x age (*t, p*) | BDI x sex (*t, p*) | BAI x age (*t, p*) | BAI x sex (*t, p*) |
|---|---|---|---|---|---|---|
| IC0 | 1.20 (0.664) | 1.37 (0.774) | 0.57 (0.898) | 1.46 (0.786) | 0.02 (0.988) | 1.10 (0.85) |
| IC1 | -0.10 (0.951) | -0.30 (0.952) | -1.04 (0.863) | -1.12 (0.786) | 0.11 (0.986) | -0.78 (0.965) |
| IC2 | 1.28 (0.664) | -0.66 (0.854) | 0.3 (0.902) | -1.37 (0.786) | 0.85 (0.914) | -0.21 (0.965) |
| IC3 | -0.80 (0.791) | 0.57 (0.854) | 0.06 (0.978) | -0.18 (0.975) | -1.72 (0.754) | -1.09 (0.85) |
| IC4 | 1.38 (0.664) | 0.85 (0.854) | 0.83 (0.884) | -0.01 (0.993) | 0.97 (0.914) | -0.85 (0.965) |
| IC8 | 0.43 (0.914) | 0.31 (0.952) | -0.40 (0.902) | 0.70 (0.799) | -0.25 (0.986) | 0.29 (0.965) |
| IC15 | -0.76 (0.791) | -1.70 (0.665) | 0.57 (0.898) | -1.11 (0.786) | 1.04 (0.914) | 1.18 (0.85) |
| IC16 | -0.43 (0.914) | -0.28 (0.952) | 0.43 (0.902) | 0.12 (0.993) | 0.78 (0.914) | 0.36 (0.965) |
| IC17 | 0.14 (0.951) | -0.74 (0.854) | 1.18 (0.863) | -0.50 (0.799) | 0.18 (0.986) | -0.03 (0.986) |
| IC20 | 0.20 (0.951) | 0.97 (0.854) | 0.46 (0.902) | 0.49 (0.799) | -0.16 (0.986) | -0.57 (0.965) |
| IC22 | 1.40 (0.664) | 0.10 (0.961) | 0.94 (0.863) | 0.06 (0.993) | 0.71 (0.914) | 0.35 (0.965) |
| IC23 | 1.18 (0.664) | 0.61 (0.854) | 0.11 (0.955) | -0.07 (0.993) | 0.44 (0.94) | 0.12 (0.978) |
| IC25 | 0.12 (0.951) | -0.69 (0.854) | -0.50 (0.902) | 0.74 (0.799) | -0.54 (0.937) | -1.09 (0.85) |
| IC26 | -0.53 (0.911) | 0.88 (0.854) | 0.87 (0.884) | 0.78 (0.799) | 0.15 (0.986) | 0.58 (0.965) |
| IC27 | 1.41 (0.664) | -0.49 (0.854) | -0.02 (0.988) | -1.78 (0.786) | -0.74 (0.914) | -2.59 (0.559) |
| IC28 | 0.50 (0.911) | -0.91 (0.854) | -1.62 (0.794) | -1.49 (0.786) | -1.53 (0.846) | -2.16 (0.559) |
| IC29 | 1.49 (0.664) | -0.14 (0.961) | 1.48 (0.802) | -1.37 (0.786) | 0.38 (0.986) | -1.52 (0.85) |
| IC30 | 0.13 (0.951) | 1.66 (0.665) | 0.76 (0.896) | 0.78 (0.799) | 0.79 (0.914) | 1.20 (0.85) |
| IC31 | 1.35 (0.664) | 0.89 (0.854) | 2.18 (0.51) | 1.43 (0.786) | 1.98 (0.754) | 0.97 (0.908) |
| IC32 | 1.67 (0.664) | 0.71 (0.854) | 2.42 (0.51) | 1.08 (0.786) | 2.22 (0.754) | 0.33 (0.965) |
| IC33 | -0.81 (0.791) | 0.03 (0.974) | -1.23 (0.863) | -1.41 (0.786) | -1.92 (0.754) | -1.46 (0.85) |
| IC34 | -1.43 (0.664) | -0.11 (0.961) | -0.96 (0.863) | -1.87 (0.786) | -0.22 (0.986) | -0.13 (0.978) |
| IC35 | 1.28 (0.664) | -0.07 (0.961) | 2.32 (0.51) | -0.32 (0.884) | 1.77 (0.754) | 0.21 (0.965) |
| IC36 | 2.11 (0.608) | 2.46 (0.425) | 1.00 (0.863) | 0.10 (0.993) | 0.84 (0.914) | 0.40 (0.965) |
| IC37 | 0.49 (0.911) | -0.11 (0.961) | 0.56 (0.898) | -1.27 (0.786) | -0.24 (0.986) | -1.63 (0.85) |
| IC38 | -0.35 (0.919) | -1.14 (0.774) | -1.43 (0.802) | -1.86 (0.786) | -1.27 (0.914) | -2.15 (0.559) |
| IC39 | 0.70 (0.793) | 0.31 (0.952) | 0.71 (0.898) | 1.20 (0.786) | 1.06 (0.914) | 0.24 (0.965) |
| IC40 | 1.01 (0.781) | 0.83 (0.854) | 1.10 (0.863) | 1.12 (0.786) | 1.65 (0.754) | 1.37 (0.85) |
| IC41 | 1.43 (0.664) | 3.18 (0.111) | 1.13 (0.863) | 0.25 (0.926) | 0.49 (0.937) | -0.65 (0.965) |
| IC42 | -0.23 (0.951) | 0.51 (0.854) | 0.63 (0.898) | -0.44 (0.993) | 0.65 (0.914) | -0.28 (0.965) |
| IC43 | -1.52 (0.664) | -0.50 (0.854) | 0.95 (0.863) | 1.42 (0.786) | 0.74 (0.914) | 0.96 (0.908) |
| IC44 | 1.35 (0.664) | 0.94 (0.854) | 1.66 (0.794) | 0.96 (0.799) | -0.09 (0.986) | -0.37 (0.965) |
| IC45 | -0.74 (0.791) | 1.71 (0.665) | 0.54 (0.898) | 0.74 (0.799) | 0.13 (0.986) | 0.23 (0.965) |
| IC46 | -1.33 (0.664) | 0.66 (0.854) | -1.02 (0.863) | -0.41 (0.827) | -0.61 (0.914) | -0.32 (0.965) |
| IC47 | 2.56 (0.374) | 0.82 (0.854) | 2.27 (0.51) | -0.36 (0.865) | 1.31 (0.914) | -0.40 (0.965) |
| IC48 | -0.38 (0.919) | -0.25 (0.961) | 0.32 (0.902) | 1.41 (0.786) | -0.08 (0.986) | 0.60 (0.965) |
| IC49 | -0.21 (0.951) | 0.17 (0.961) | -1.77 (0.794) | -1.04 (0.799) | -1.96 (0.754) | -1.23 (0.85) |
| IC50 | 1.66 (0.664) | 0.51 (0.854) | -0.23 (0.902) | -0.51 (0.799) | -0.46 (0.94) | -1.26 (0.85) |
| IC51 | 0.36 (0.919) | -1.48 (0.774) | 0.23 (0.902) | -0.96 (0.799) | 0.06 (0.986) | -1.18 (0.85) |
| IC52 | 1.18 (0.664) | 0.55 (0.854) | 0.23 (0.902) | -0.50 (0.799) | 0.07 (0.986) | 0.02 (0.986) |
| IC53 | 0.76 (0.791) | 0.13 (0.961) | 0.69 (0.898) | 1.081 (0.786) | -0.26 (0.986) | -0.55 (0.965) |
| IC54 | -0.08 (0.953) | 0.29 (0.952) | -0.37 (0.902) | -0.69 (0.799) | -0.80 (0.914) | -1.00 (0.908) |
| IC55 | -0.01 (0.995) | -0.65 (0.854) | 1.42 (0.802) | -0.83 (0.799) | 1.20 (0.914) | 0.31 (0.965) |
| IC56 | -3.16 (0.118) | -0.61 (0.854) | -2.07 (0.535) | -0.92 (0.799) | -2.36 (0.754) | -2.14 (0.559) |
| IC57 | -1.26 (0.664) | 1.35 (0.774) | -0.59 (0.898) | 0.77 (0.799) | -0.51 (0.937) | 0.07 (0.986) |
| IC58 | -0.81 (0.791) | -0.80 (0.854) | -1.24 (0.863) | -1.44 (0.786) | -0.74 (0.914) | -1.27 (0.85) |
| IC59 | 0.91 (0.781) | -2.09 (0.425) | 1.57 (0.797) | -1.13 (0.786) | 0.93 (0.914) | -0.26 (0.965) |
| IC60 | 0.37 (0.919) | 2.18 (0.425) | 0.46 (0.902) | 0.65 (0.799) | 0.52 (0.937) | 1.27 (0.85) |
| IC61 | -0.89 (0.781) | 1.81 (0.665) | 0.36 (0.902) | -0.02 (0.993) | 0.03 (0.988) | 0.31 (0.965) |
| IC62 | 0.29 (0.921) | 0.11 (0.961) | 0.91 (0.873) | -0.43 (0.827) | 1.08 (0.914) | -0.19 (0.965) |
| IC63 | 0.91 (0.781) | 0.29 (0.952) | 0.28 (0.902) | -0.53 (0.799) | 0.61 (0.914) | -1.16 (0.85) |
| IC64 | -1.00 (0.781) | -2.27 (0.425) | -0.03 (0.988) | -1.09 (0.786) | 0.12 (0.986) | -0.76 (0.965) |
| IC65 | 0.32 (0.921) | 1.20 (0.774) | 0.55 (0.898) | 1.57 (0.786) | 0.29 (0.986) | 0.56 (0.965) |
| IC66 | 0.30 (0.921) | -0.77 (0.854) | -0.64 (0.898) | -2.01 (0.786) | -1.47 (0.875) | -1.42 (0.85) |
| IC67 | 2.16 (0.608) | 1.15 (0.774) | -1.28 (0.863) | -0.52 (0.799) | -1.29 (0.914) | 0.20 (0.965) |
| IC68 | -0.70 (0.793) | 0.08 (0.961) | 0.28 (0.902) | 0.04 (0.993) | 0.72 (0.914) | -0.29 (0.965) |
| IC69 | 1.04 (0.781) | 1.23 (0.774) | 0.37 (0.902) | -0.57 (0.799) | -0.35 (0.986) | -1.32 (0.85) |
| IC70 | -0.79 (0.791) | -1.18 (0.774) | -0.80 (0.894) | -0.66 (0.799) | -0.63 (0.914) | -0.25 (0.965) |

| | | | | | | |
|---|---|---|---|---|---|---|
| IC71 | 0.17 (0.951) | -1.17 (0.774) | 0.31 (0.902) | 0.79 (0.799) | 0.92 (0.914) | 1.85 (0.85) |
| IC72 | 0.10 (0.951) | -0.53 (0.854) | 0.11 (0.955) | -0.48 (0.799) | 0.06 (0.986) | 0.02 (0.986) |
| IC73 | -0.53 (0.911) | -1.55 (0.748) | -0.31 (0.902) | -0.89 (0.799) | -0.80 (0.914) | 0.02 (0.986) |
| IC74 | -1.72 (0.664) | -2.09 (0.425) | -0.85 (0.884) | 0.51 (0.799) | -0.64 (0.914) | 0.51 (0.965) |
| IC75 | -0.93 (0.781) | 0.19 (0.961) | -0.75 (0.896) | 0.78 (0.799) | -0.92 (0.914) | 0.14 (0.978) |
| IC76 | 0.45 (0.914) | -1.29 (0.774) | -1.63 (0.794) | -0.90 (0.799) | -0.69 (0.914) | 0.32 (0.965) |
| IC77 | -1.18 (0.664) | -0.74 (0.854) | -1.12 (0.863) | 0.49 (0.799) | -1.67 (0.754) | 0.47 (0.965) |
| IC78 | -0.64 (0.839) | -1.20 (0.774) | -0.95 (0.863) | -1.09 (0.786) | -0.62 (0.914) | -0.48 (0.965) |
| IC79 | -0.95 (0.781) | -1.00 (0.854) | -0.16 (0.941) | -0.60 (0.799) | 0.48 (0.937) | -0.60 (0.965) |

### Figures

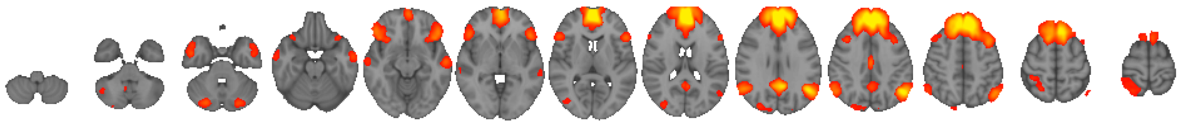

**Fig. S1.** DMN resting-state network extracted from the rs-fMRI data using FSL MELODIC

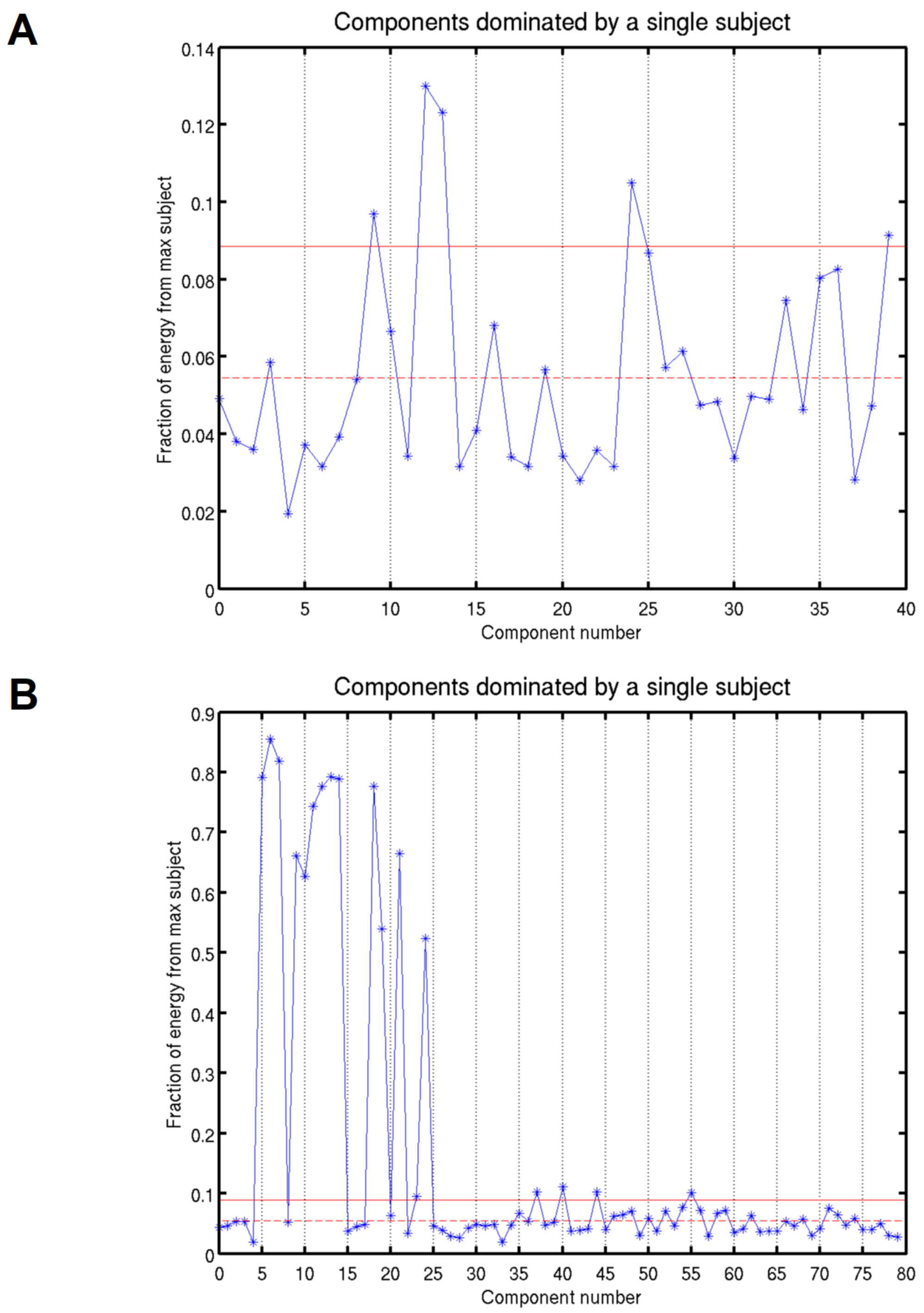


**Fig. S2**. Lineplots showing the degree to which a component is dominated by a single subject based on the fraction of energy in the decomposition with (A) 40 components (main analyses) and (B) 80 components (supplementary analyses).

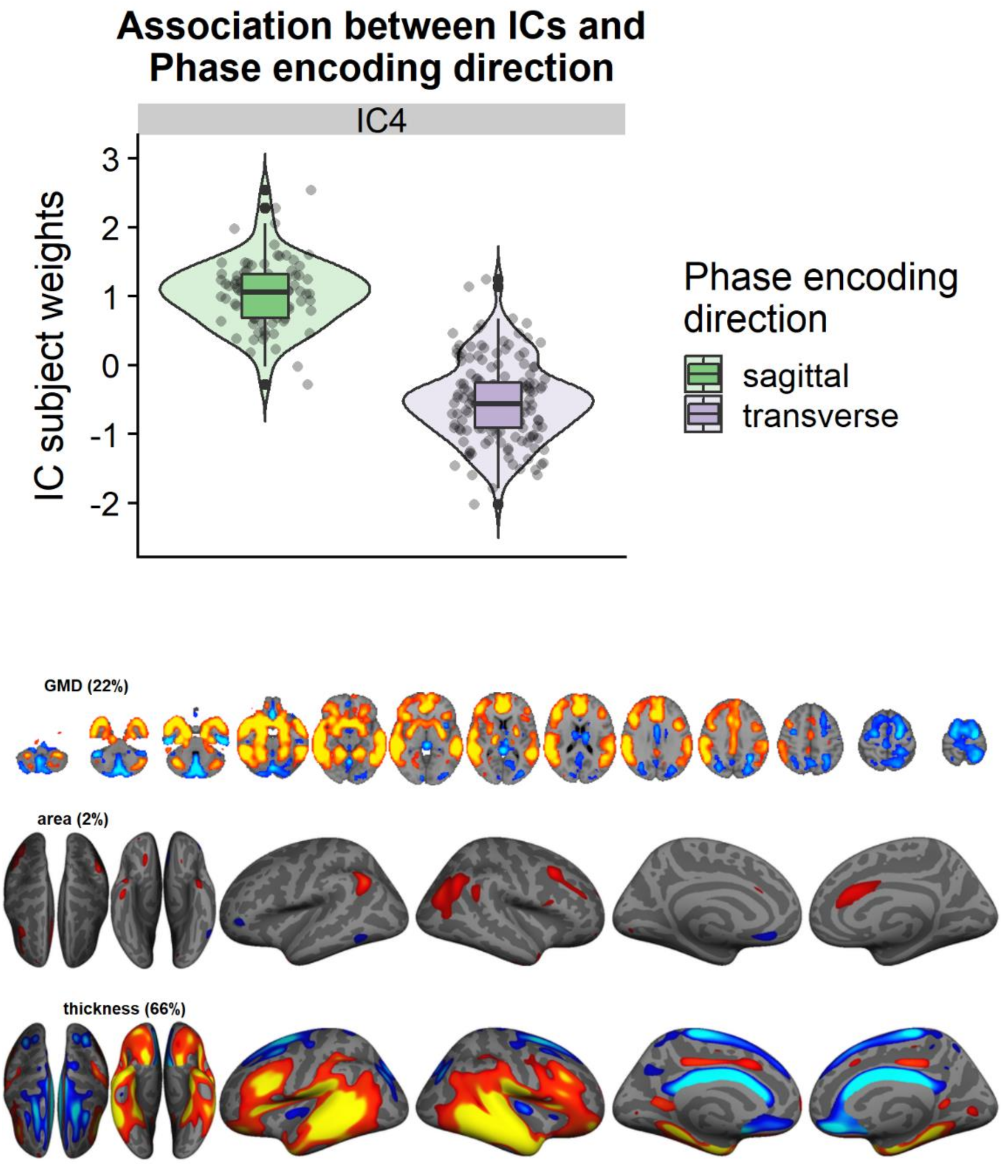


**Fig. S3.** The LICA component that was very sensitive to phase encoding direction. The subject weights have been residualized for group, age and sex.

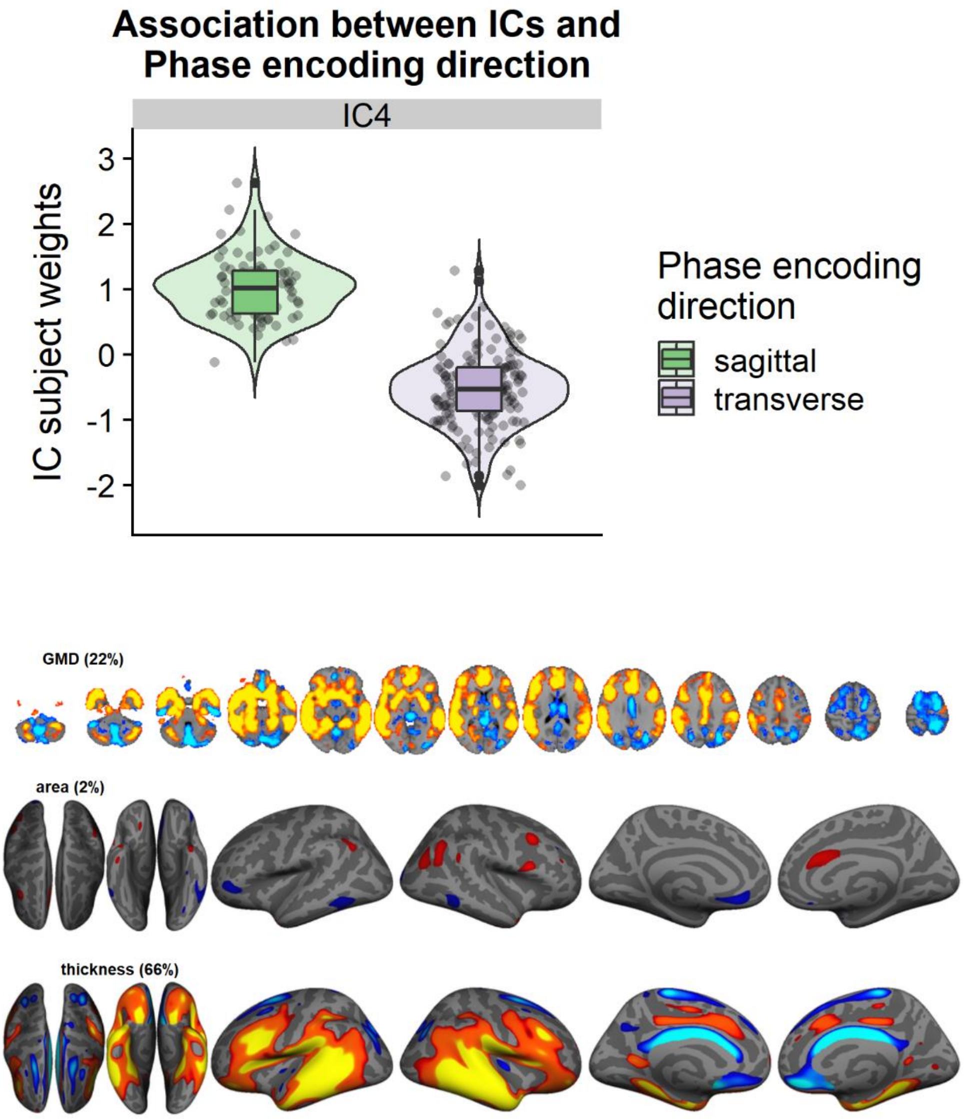


**Fig. S4.** The LICA component that was very sensitive to phase encoding direction in the higher decomposition. The subject weights have been residualized for group, age and sex.

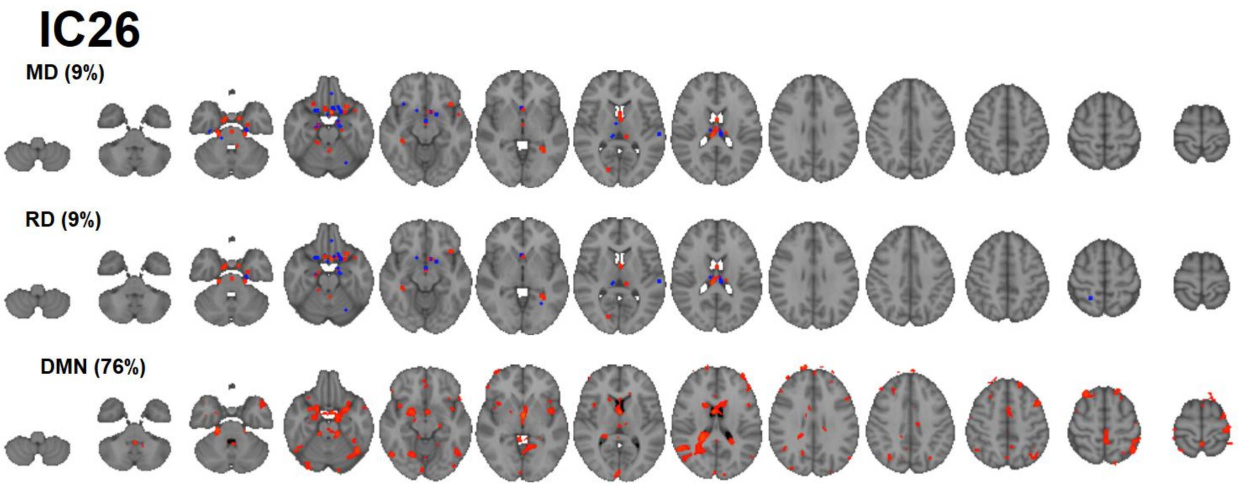


**Fig. S5.** The other component that had substantial contribution from resting-state DMN FC.

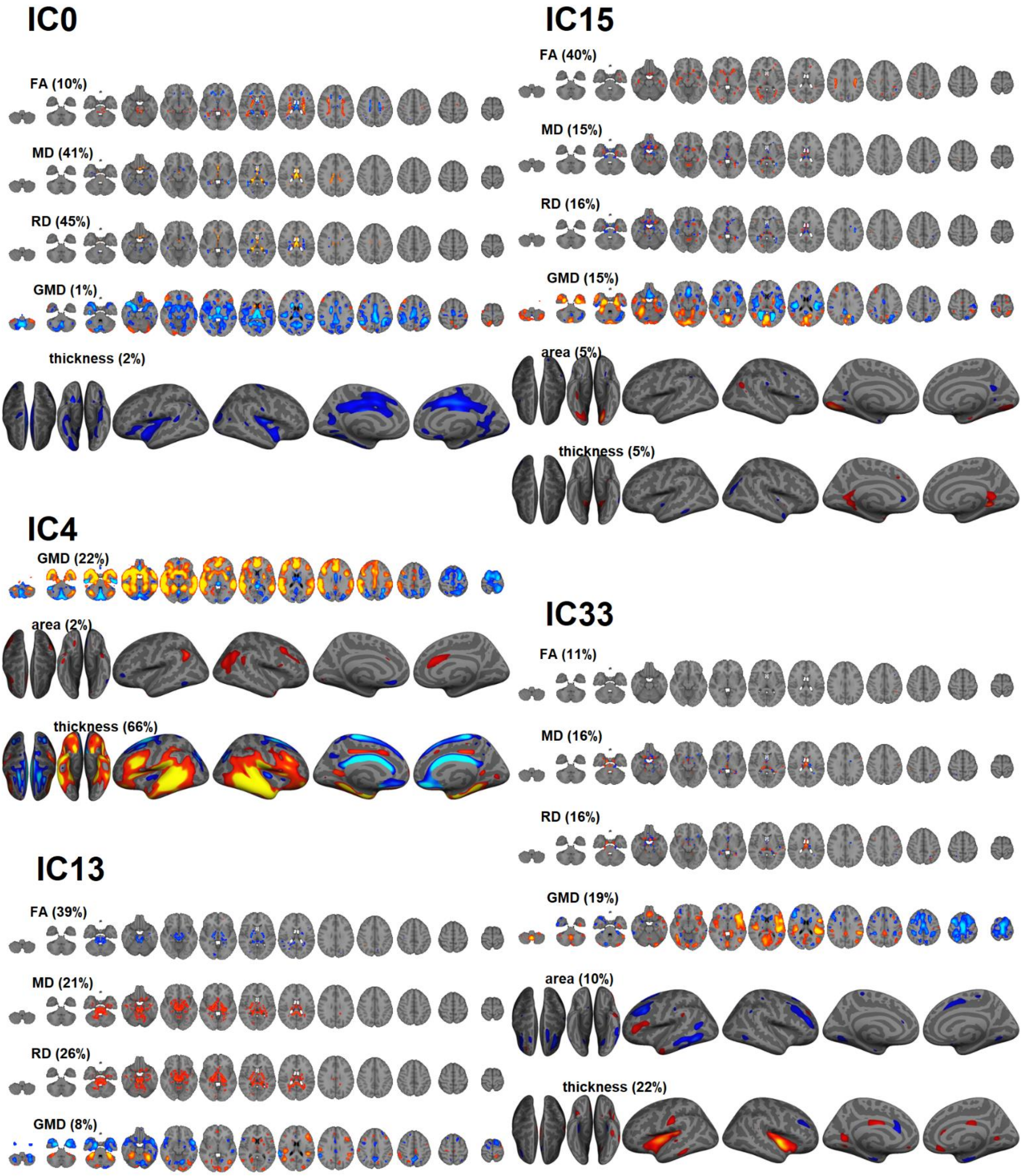


**Fig S6.** ICs characterized by region specific features that are associated with age or sex.

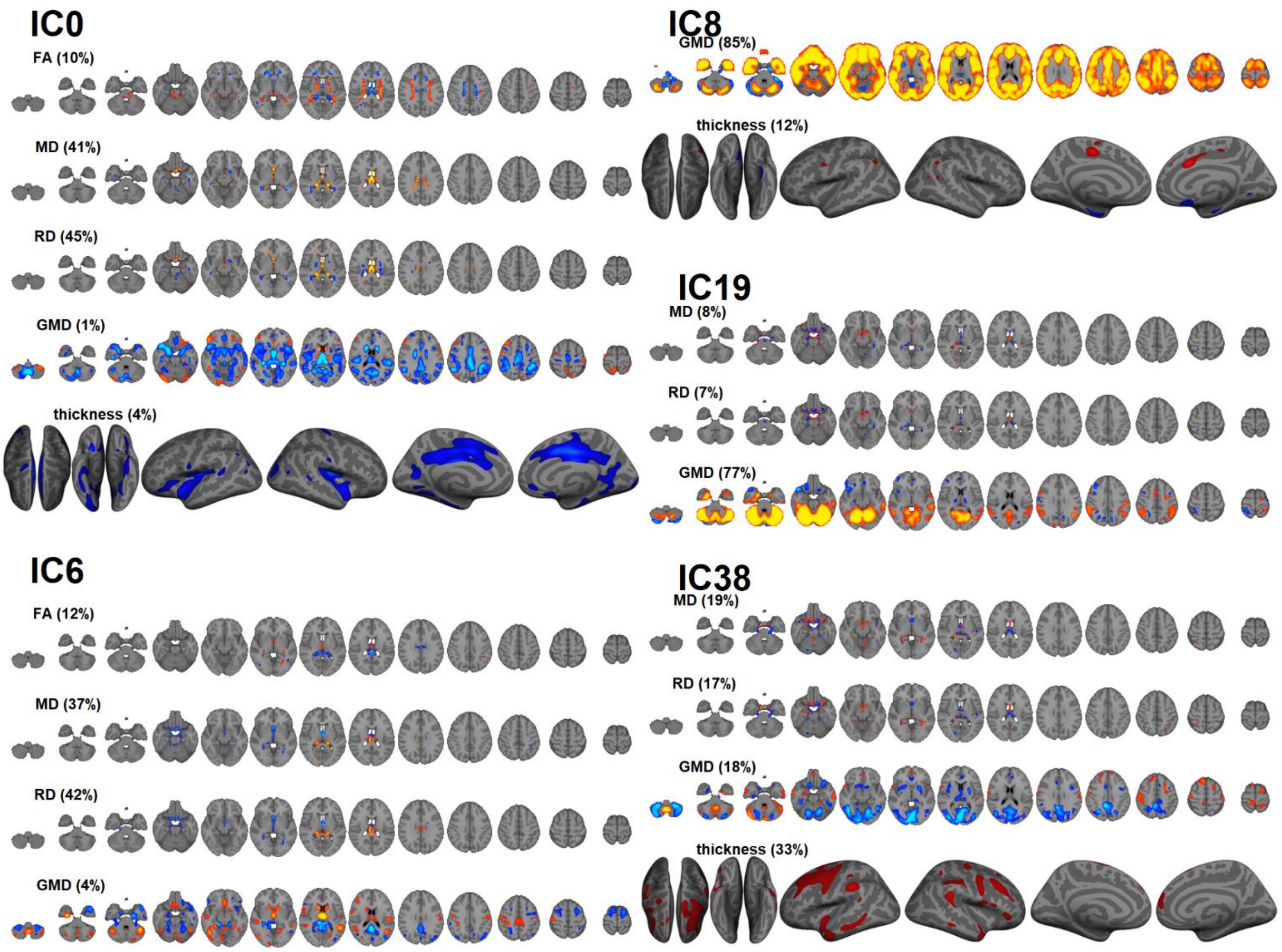


**Fig. S7.** Some of the top features (ICs) in the machine learning analyses. IC0 is among the top features for classifying case-control status and predicting symptom load for depression. IC6 is among the top features for predicting symptom loads for depression and anxiety. IC8 is among the top features for classifying case-control status and predicting symptom load for anxiety. IC19 is the top feature for classifying case-control status. IC38 is among the top features for predicting symptom loads for depression and anxiety.

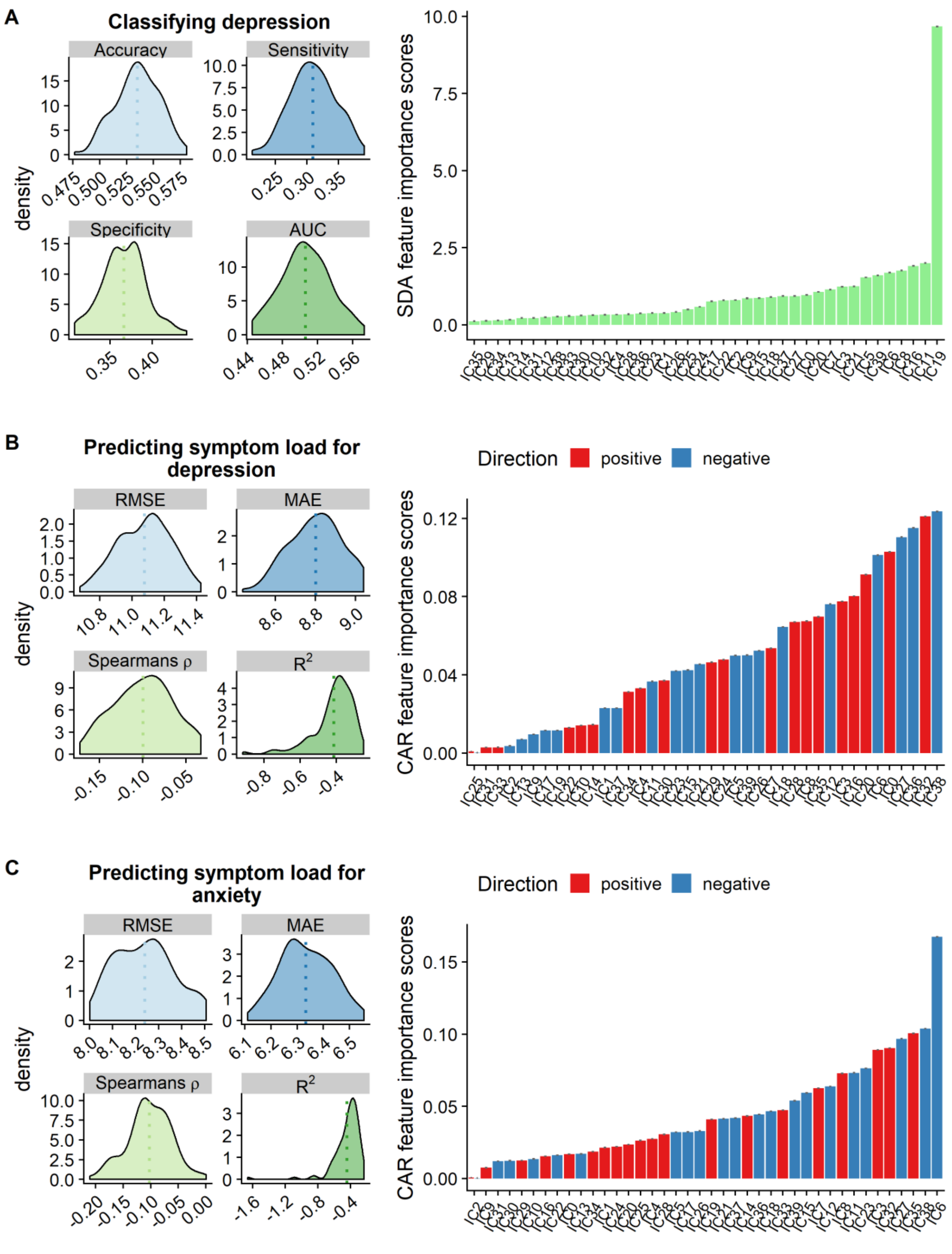


**Fig. S8.** The results of the supplementary analyses of the machine learning approach using 10-fold cross-validation with 100 repetitions for (A) classifying case-control status (B) prediction symptom load for depression and (C) symptom load for anxiety. Here, age, sex and phase encoding were regressed out from the subject weights of all the ICs. The figures on the left show prediction accuracy based on various model performance metrics. The barplots on the right show the most important features for each model based on CAT-scores (A) or CAR-scores (B and C).

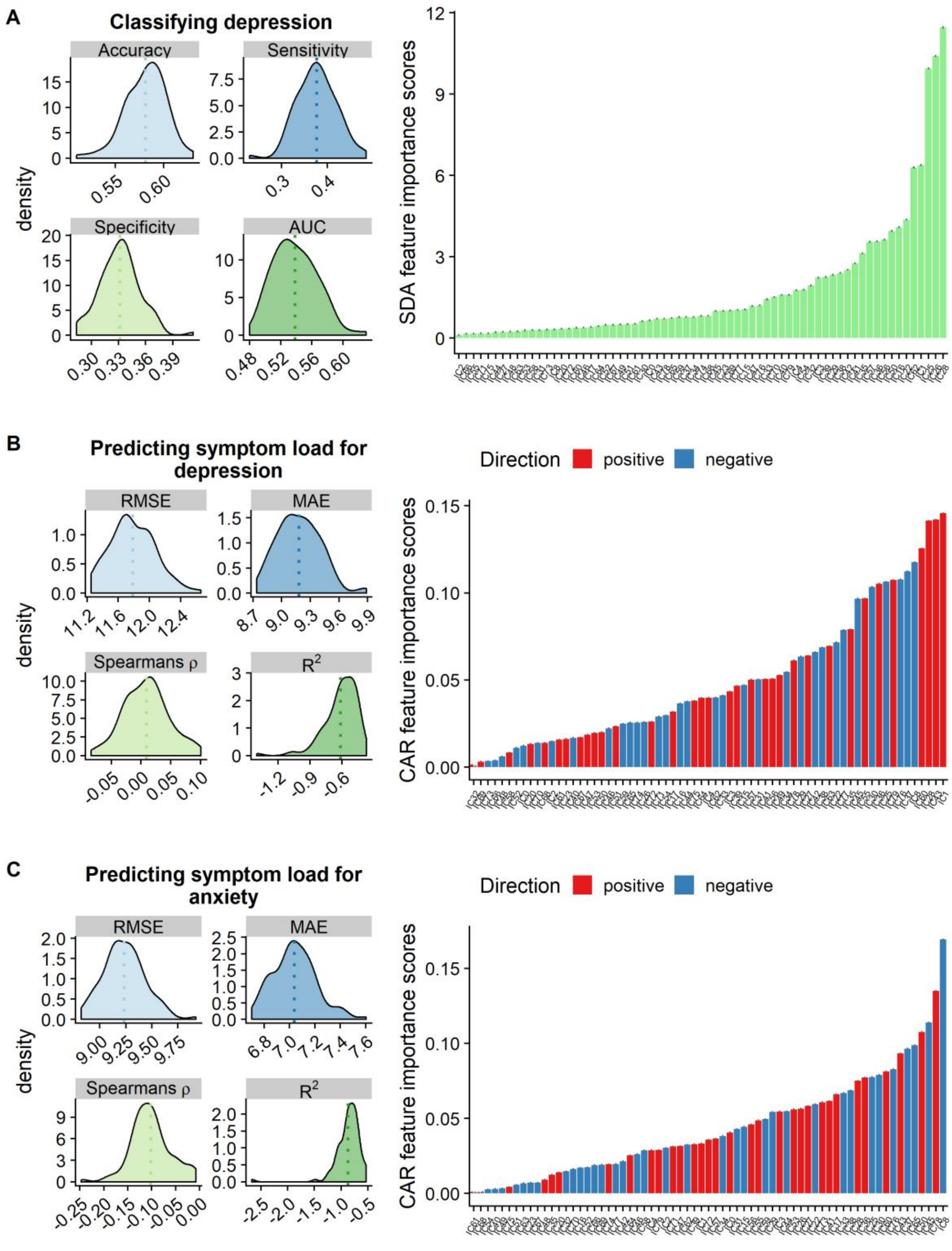


**Fig. S9.** The results of the machine learning approach based on the 80 IC decomposition (67 ICs in total) using 10-fold cross-validation with 100 repetitions for (A) classifying case-control status (B) prediction symptom load for depression and (C) symptom load for anxiety. Here, phase encoding direction was only regressed out of the subject weights in IC4, while age and sex were regressed out from the subject weights of all the ICs. The figures on the left show prediction accuracy based on various model performance metrics. The barplots on the right show the most important features for each model based on CAT-scores (A) or CAR-scores (B and C).

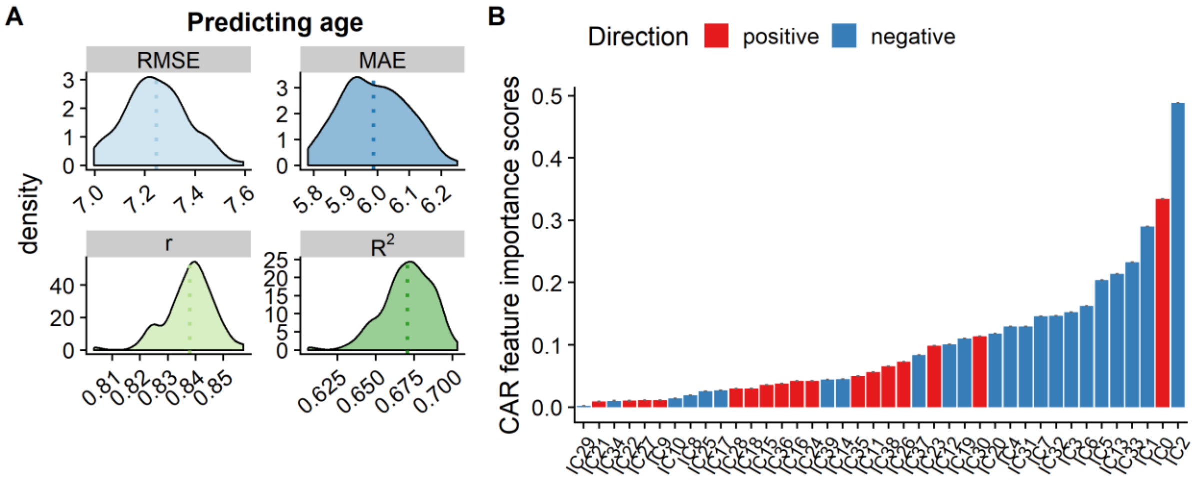


**Fig. S10.** Age prediction regressing out phase from the subject weights of all the ICs. (A) model performance results and (B) feature importance

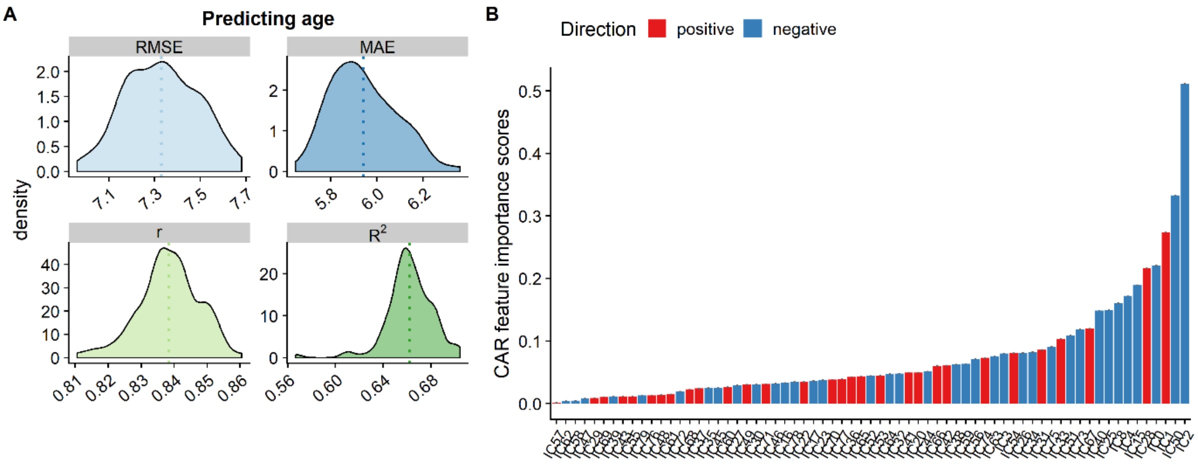


**Fig. S11**. Age prediction based on the 80 IC decomposition (67 ICs in total), regressing out phase encoding from the subject weights in IC4. (A) model performance results and (B) feature importance
